# How do species barriers decay? Concordance and local introgression in mosaic hybrid zones of mussels

**DOI:** 10.1101/818559

**Authors:** Alexis Simon, Christelle Fraïsse, Tahani El Ayari, Cathy Liautard-Haag, Petr Strelkov, John J. Welch, Nicolas Bierne

## Abstract

The *Mytilus* complex of marine mussel species forms a mosaic of hybrid zones, found across temperate regions of the globe. This allows us to study “replicated” instances of secondary contact between closely-related species. Previous work on this complex has shown that local introgression is both widespread and highly heterogeneous, and has identified SNPs that are outliers of differentiation between lineages. Here, we developed an ancestry-informative panel of such SNPs. We then compared their frequencies in newly-sampled populations, including samples from within the hybrid zones, and parental populations at different distances from the contact. Results show that close to the hybrid zones, some outlier loci are near to fixation for the heterospecific allele, suggesting enhanced local introgression, or the local sweep of a shared ancestral allele. Conversely, genomic cline analyses, treating local parental populations as the reference, reveal a globally high concordance among loci, albeit with a few signals of asymmetric introgression. Enhanced local introgression at specific loci is consistent with the early transfer of adaptive variants after contact, possibly including asymmetric bistable variants (Dobzhansky-Muller incompatibilities), or haplotypes loaded with fewer deleterious mutations. Having escaped one barrier, however, these variants can be trapped or delayed at the next barrier, confining the introgression locally. These results shed light on the decay of species barriers during phases of contact.

## 1 Introduction

Divergence between species is almost always accompanied by hybridisation during phases of contact (Abbott et al., 2013), and so the study of speciation is intertwined with the study of introgression (Aeschbacher, Selby, Willis, & Coop, 2017; Chaturvedi et al., 2019; Duranton et al., 2018; Gagnaire et al., 2018; Martin, Davey, Salazar, & Jiggins, 2019; Roesti, Moser, & Berner, 2013; Schumer et al., 2018). The outcome of any given contact will depend on intrinsic and extrinsic factors. When barriers are weak there can be massive introgression, leaving differentiation at only a few strongly selected loci. But even strong barriers can be rapidly crossed by adaptive alleles and flanking neutral variants (Barton, 1979a; Faure, David, Bonhomme, & Bierne, 2008).

If some variation is bi-stable (Barton & Turelli, 2011) such that one parental genotype is fitter, but heterozygous or recombinant genotypes are unfit, the spread of the fittest genotype is hindered at the barrier, and thereby contributes to the barrier (Barton, 1979a; Barton & Bengtsson, 1986; Piálek & Barton, 1997). Stochastic processes, such as random drift or variable migration rates, can free the spreading wave of these variants one by one (Piálek & Barton, 1997). The overall result is a sequential decay of the species barrier. However, the process can be very slow, and so difficult to observe in action.

A promising system to study these processes is the *Mytilus* complex of marine mussels, comprising the three species *M. trossulus*, *M. edulis* and *M. galloprovincialis*. Current transoceanic isolation (e.g. East and West Atlantic in *M. edulis* and *M. trossulus*, Riginos and Henzler, 2008; Varvio, Koehn, and Väinolä, 1988) and ancient allopatry (e.g. Atlantic and Mediterranean lineages in *M. galloprovincialis*, El Ayari, Trigui El Menif, Hamer, Cahill, and Bierne, 2019; Quesada, Zapata, and Alvarez, 1995) subdivides these taxa into infra-specific lineages (Fraïsse, Belkhir, Welch, & Bierne, 2016; Simon et al., 2020). However, they are broadcast spawners with a dispersive larval phase, and so true geographic isolation is relatively rare. For this reason, the complex is divided by numerous post-glacial hybrid zones, maintained by both intrinsic and extrinsic mechanisms of reproductive isolation (Bierne, Borsa, et al., 2003; El Ayari et al., 2019; Fraïsse, Belkhir, et al., 2016; Hilbish et al., 2012; Riginos & Cunningham, 2005; Strelkov, Katolikova, & Väinolä, 2017). In particular, in the northern hemisphere all three species form a geographic mosaic delimited by many hybrid zones (Fraïsse, Belkhir, et al., 2016).

This biographical history has two important consequences. First, contacts between similar lineages occur multiple times, and so we can observe the same species barrier in multiple demographic and ecological contexts (Abbott et al., 2013; Harrison & Larson, 2016; Simon et al., 2020). Second, with its narrow hybrid zones, combined with very long-range admixture (Fraïsse, Belkhir, et al., 2016), this complex provides us with a continuum of divergence between the interacting taxa. Precisely because of this heterogeneity, however, it is challenging to identify diagnostic loci for the *Mytilus* complex.

Recently, Fraïsse, Belkhir, et al. (2016) captured 1269 contigs of a few kb each from reference populations of the *Mytilus* complex. Using these data, they identified regions that were divergent between species, and outliers of differentiation between populations of the same species. These intraspecific outliers are significant in two ways. First, they can be used to identify individual lineages. Second, phylogenetic analyses of gene trees suggested that the majority of the outliers were due to local introgression between species, rather than local intra-specific selective sweeps. As such, they provide information about introgression in the complex.

Here, we extend the analyses of (Fraïsse, Belkhir, et al., 2016). First, we built a SNP panel of a few hundred loci chosen from their outlier contigs, and therefore enriched in highly differentiated markers. Having included intra-specific outlier loci, we then examined their allele frequencies in a wider range of populations, allowing us to examine local introgression on a broader scale. Finally, we also sampled within hybrid zones, and carried out genomic cline / concordance analyses (Gompert & Buerkle, 2011; Macholán et al., 2011). Our aim here was to test the effect of the choice of parental populations (local *versus* distant) in the detection of discordant behavior.

After verifying that our SNP panel is effective in identifying the lineages of interest, our results confirm that local introgression is both widespread in these *Mytilus* lineages, and heterogeneous across the genome. At some loci identified as intra-species outliers in (Fraïsse, Belkhir, et al., 2016), heterospecific alleles are sometimes fixed, or nearly so, in one parental patch close to a hybrid zone, while remaining nearly absent from other population patches farther from the zone. Genomic clines suggest high concordance among loci in the centre of hybrid zones, but only when local parental populations are used as reference. However, a few loci do depart from the genomic average, and demonstrate moderate asymmetric introgression. Unlike outliers that exhibit enhanced local introgression at a large scale, the introgression of these genomic cline outliers does not extend outside of the hybrid zones where lineages coexist.

## 2 Materials and methods

### 2.1 Sampling

*Mytilus spp.* individuals were sampled from 58 locations, including several known hybrid zones (Figure 1, Table 1). Sampling sites are located on the American Pacific coast, the American and European North Atlantic coasts, and in the Mediterranean, Baltic, North, Barents and Black seas. 441 individuals were newly genotyped and 72 genotypes by sequencing (GBS) were extracted from the published sequence dataset of Fraïsse, Belkhir, et al. (2016).

**Table 1:** Definition of reference populations. The ID column gives the population number referenced in Figure 1 and Table S1. Abbreviations used in the manuscript and figures are combinations of text in parentheses (e.g., edu_eu_north).

| Level 1 | Level 2 | Level 3 | comment | ID |
| --- | --- | --- | --- | --- |
| *M. trossulus* (tros) | America (am) | - | Saint Lawrence | 08, 09 |
|  | Europe (eu) | North (north) | Russia and Scandinavia | 18, 19 |
|  |  | Baltic (baltic) | Baltic sea | 22 |
|  | Pacific (pac) | East (east) | From Californian coast | 01, 02, 05 |
| *M. edulis* (edu) | America (am) | - | Long Island region | 10-17 |
|  | Europe (eu) | external (ext) | - | 26-28 |
|  |  | local (int) | - | 35-37 |
|  |  | North (north) | Russia and Scandinavia | 18, 19 |
| *M. galloprovincialis* (gallo) | Atlantic (atl) | external (ext) | - | 40-41 |
|  |  | local (int) | - | 31-32 |
|  | Mediterranean (med) | West (west) | - | 51-54 |
|  |  | East (east) | - | 55-57 |
|  |  | Black Sea (bs) | - | 58 |

**Figure 1:**
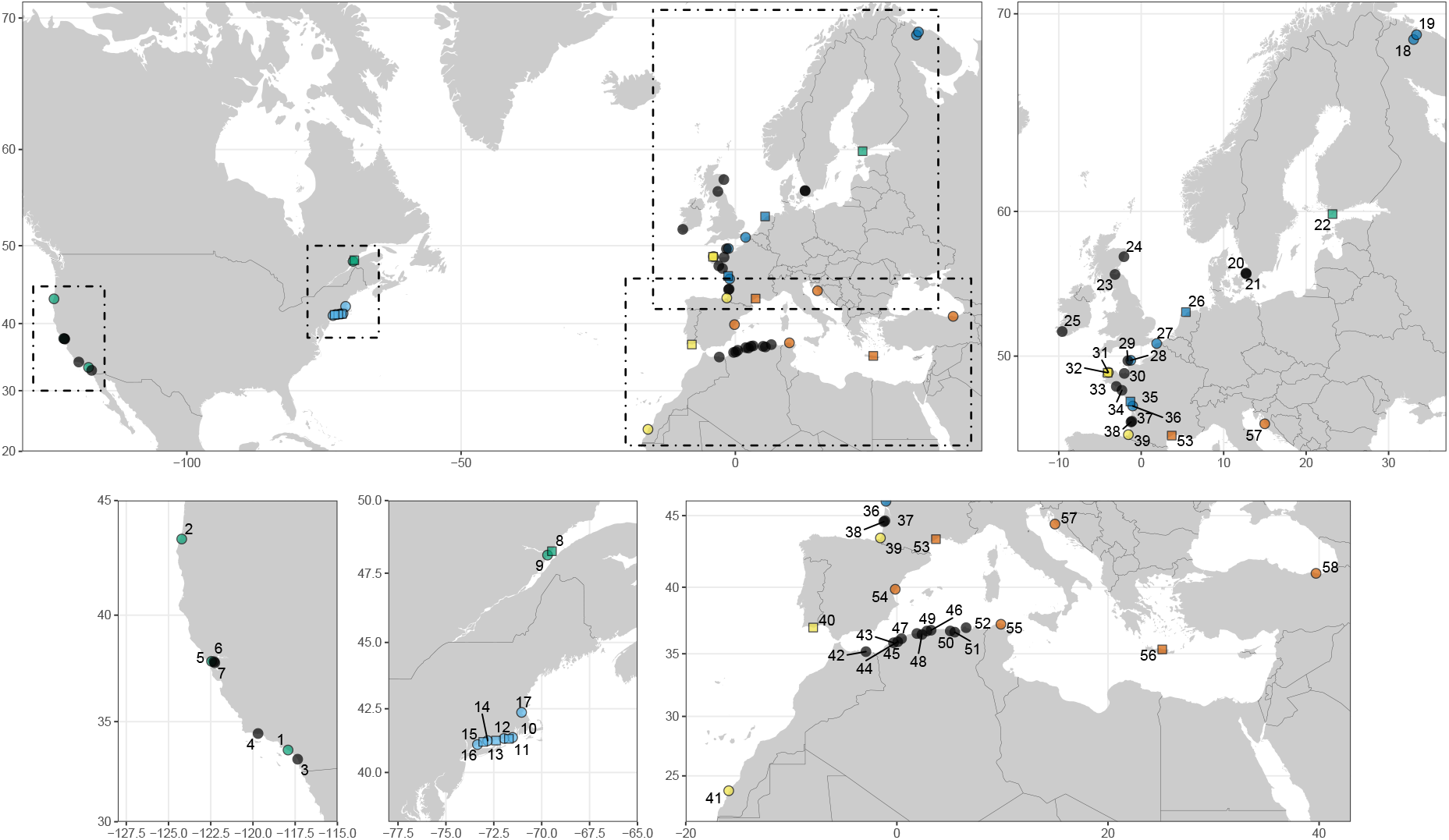
Location of the studied populations. Squares: genotype by sequencing (GBS, data from Fraïsse, Belkhir, Welch, and Bierne, 2016). Circles: BeadXpress genotypes. More detailed population information is available in Table S1.

### 2.2 Assay design

We aimed to genotype previously identified differentiated and outlier loci, across a large number of samples, in a cost-effective manner. For this purpose we used an Illumina BeadXpress® assay with Veracode™ technology (GoldenGate Genotyping Assay). This method is based on the hybridisation and PCR amplification of SNP- and locus-specific oligos differentially bound to two different dies. The genotype of an individual is provided by the relative fluorescence. We designed an assay of 384 SNPs (being the multiplexing limit of the technology).

Loci were selected, prior to genotyping, based on their differentiation either between species or between populations of the same species, using the published results of (Fraïsse, Belkhir, et al., 2016). Briefly, this database was produced via a target enrichment method on the three species and multiple populations of the *Mytilus* species complex (Fraïsse, Belkhir, et al., 2016). It contains 1269 contigs sequenced for 72 individuals from eleven different locations http://www.scbi.uma.es/mytilus/index.php.

Markers with a minimum allele frequency of 0.05 and a maximum missing data percentage of 50% computed on all individuals were retained. Coverage was estimated as a mean computed on three populations (two Atlantic *M. galloprovincialis* and one Atlantic *M. edulis*). Contig regions with especially high coverage (> 300 reads) were excluded to avoid repeated elements. Regions of the database produced from cDNA were blasted against a draft genome (Murgarella et al., 2016) to exclude regions close to intron/exon limits as flanking sequences in 3’ and 5’ of the target SNP were needed to design primers for PCR amplification. For the same reason, SNPs close to the start and end of the contigs were also excluded. An ADT score, produced by Illumina Assay Design Tool, quantifies the expected amplification success, and was used to filter the SNPs with the most probable design success (ADT score > 0.4). *F*_*ST*_ was computed between pairs of populations and species such as those shown in Table 2. A first filtration step kept the 5 highest *F*_*ST*_ markers per contig for each comparison. We ensured a *F*_*ST*_ > 0.5 for all comparisons, except within edu_am and between gallo_atl and gallo_med where we used a *F*_*ST*_ > 0.4. Then, the six SNPs with the highest ADT scores per contig were retained. Finally, one SNP per contig and per category with the highest *F*_*ST*_ was selected (Table 2). Only for the *M. trossulus* comparison, the 70 SNPs with the highest ADT scores were retained. This SNP selection provides a final dataset of 384 SNPs, balanced across comparisons (Table 2).

**Table 2:** Number of SNPs per type of comparison. The SNP added to the dataset (extra) corresponds to an introgressed amino-acid changing variant (see main text). “all” corresponds to the other populations.

| Comparison | Number of SNPs genotyped | Number of SNPs retained |
| --- | --- | --- |
| edu / all | 2 | 2 |
| edu_am / edu_eu | 39 | 22 |
| within edu_am | 17 | 5 |
| edu_eu / all | 51 | 16 |
| gallo_eu / all | 12 | 4 |
| gallo_atl / gallo_med | 27 | 17 |
| gallo / edu | 100 | 65 |
| gallo_med / all | 6 | 1 |
| tros_am / tros_eu | 34 | 12 |
| tros / all | 96 | 68 |
| Total | 384 | 212 |

### 2.3 Genotyping and filtering

Genotyping of the 441 individuals was performed with the BeadXpress® (hereafter BXP) technology.

Among the 384 SNPs genotyped, 252 were readable (clusters of homozygotes/heterozygotes well defined) and 132 were lost. This low rate of successful amplification is expected in such highly polymorphic species and because of our choice to retain only good quality SNPs. 40 additional SNPs were removed due to a differentiation between the BXP and GBS typing within populations. The threshold of missing data per SNP and per individual was set to 10%. One marker (190) was rescued as the missing data was mostly due to low amplification in *M. trossulus* only (overall 12% missing data).

The genotypes for the 77 GBS individuals were retrieved from the calling in Fraïsse, Belkhir, et al. (2016).

Two replicated individuals between the GBS and BeadXpress were present in the dataset.

They showed mismatch levels of, respectively, 2.83% and 2.36% between the two experiments, and this was due entirely to a well-known heterozygote assignment bias in GBS experiments. The two replicated GBS individuals were removed from further analyses.

Three samples were removed from the analyses prior to filtering as they have already proven to be cancerous individuals (Riquet, Le Cam, Fonteneau, & Viard, 2016, July 2015) (Por_40 from population 34; Arsud_05, Arsud_07 from population 38).

Reference populations were defined based on previous knowledge of the *Mytilus* species complex (Fraïsse, Gunnarsson, Roze, Bierne, & Welch, 2016; Simon et al., 2020). They are used first as a classification reference for newly genotyped individuals, to compute allele frequencies in analyses, and to serve as parental populations. We used three levels of differentiation representing the species level (L1), the ocean basin or continent (L2) and regional groupings (L3) (Table 1). We used the GBS samples from Fraïsse, Belkhir, et al. (2016) as references, and previously untyped individuals were assigned to a reference group if they belonged to the same GBS population or were geographically close from it. We used a preliminary `Admixture` analysis (v1.3.0; Alexander, Novembre, and Lange, 2009) at the species level (L1) using *K* = 3 and default settings (30 independent runs merged with `CLUMPAK`; v1.1; Kopelman, Mayzel, Jakobsson, Rosenberg, and Mayrose, 2015) to correct the reference groups for inter-specific migrants or hybrids, as sympatry and hybridisation is a common phenomenon in the *Mytilus* species complex. Individuals assigned to those populations, and not filtered out, constitute the “reference dataset”.

Due to suspicious levels of *M. trossulus* ancestry in locations devoid of this species, which could indicate the presence of a transmissible cancer (Riquet et al., 2016, July 2015), two additional samples from the Mediterranean sea were removed (MTP_05 and Collo_10 from populations 48 and 52 respectively).

A more stringent filtration was additionally applied to the `edu_am` population when used as a reference, given the presence of European ancestry in the Long Island Sound (populations 15 and 16, Figure S10, also described in Simon et al., 2020). Filtering yielded a total of 514 individuals genotyped at 212 markers.

Hardy-Weinberg equilibrium was tested for the remaining markers within putative panmictic clusters outside of known hybrid zones (`pegas` 0.11, Paradis, 2010). Exact tests were performed using 1000 bootstraps and p-values were adjusted for false discovery rate using the Benjamini-Yekutieli method (Benjamini & Yekutieli, 2001).

We used a partial genetic map produced in Simon et al. (2020) based on an F2 cross between Mediterranean *M. galloprovincialis* and *M. edulis*, genotyped at a subset of the markers studied here (Bierne, Bonhomme, Boudry, Szulkin, & David, 2006; Bierne, David, Boudry, & Bonhomme, 2002; Simon, Bierne, & Welch, 2018). In addition to this genetic map, we extrapolated genetic distances from physical distance for markers present on the same contig using a recombination rate of 2 cM/Mb (rounded from the estimate in Bierne, 2010).

### 2.4 Population structure

A principal components analysis (PCA) was performed using markers on different physical contigs (retaining the one with least missing data). This filtration was to avoid strong biases due to physical linkage, and led to a final set of 160 markers. The genotype data was centred and scaled using the `adegenet` R package (Jombart, 2008), with the replacement of missing data by the mean allele frequencies. Following the PCA, a dimensional reduction method, UMAP (Diaz-Papkovich, Anderson-Trocme, & Gravel, 2019; McInnes & Healy, 2018), was applied to the first 11 principal components. This threshold was chosen based on the expectation of 12 panmictic populations (level L3 of the reference groups, Diaz-Papkovich et al., 2019). This method was performed using the python package `umap-learn` (McInnes & Healy, 2018) and the R wrapper package `umap` (Konopka, 2019).

The `Structure` software (v2.3.4; Falush, Stephens, and Pritchard, 2003) was used to provide Bayesian estimates of ancestry with an admixture model. `Structure` was run with the admixture and linkage model (LINKAGE = 1). The dataset was filtered with the following steps: (i) markers out of Hardy-Weinberg equilibrium were removed; (ii) one marker per physical contig was retained (keeping the one with least missing data); and (iii) the genetic map was used to produce linkage information for the retained markers, with the assumption that markers absent from the genetic map were unlinked (for lack of further information).

For the global `Structure` analysis, 20 independent Monte Carlo Markov Chains (MCMC) runs of 20,000 burn-in iterations followed by 80,000 steps were performed to estimate model parameters for each *K* between 4 and 12. The standard deviation for the *α* prior was set to 0.05 for better mixing of the chains. The `Clumpak` software (Kopelman et al., 2015) was used to investigate and aggregate `Structure` outputs with an MCL threshold of 0.9. Only major clusters are presented in the results.

### 2.5 Hybrid zone analyses

Each hybrid zone was defined by parent 1 (P1), parent 2 (P2) and central populations (Table 3). Each parental population was classified as either “local” or “distant”, according to observed levels of introgression or geography. A local parental population is defined as a population close to the focal hybrid zone and being potentially subject to recent gene flow from the hybrid zone. A distant parental population is defined as a population distant from the hybrid zone, which has been less influenced by recent gene flow from it but can still have a substantial rate of established introgression due to past contacts. In a few cases a local parental population was not available in our sampling design and in these cases, a distant population was used instead (Table 3).

**Table 3:** Grouping of populations for each hybrid zone considered in this study. We consider parental populations (P1/P2) from either the distant or the local zone. See Figure 1-2 and Table S1 for population information. In the populations listed for parental groups, only the individuals representative of the focal group are retained (e.g. only *M. edulis* individuals in populations 18 and 19 in the Øresund analysis).

| Hybrid zone | Lineages | P1 distant | P1 local | Central | P2 local | P2 distant |
| --- | --- | --- | --- | --- | --- | --- |
| Øresund | edu_eu/tros_eu_baltic | 18, 19 | – | 20, 21 | 22 | 08, 09 |
| Scotland | edu_eu/gallo_atl | 26-27 | – | 24 | 31, 32 | 40, 41 |
| Brittany | edu_eu/gallo_atl | 26-27 | 35-37 | 33, 34 | 31, 32 | 40, 41 |
| Aquitaine | edu_eu/gallo_atl | 26-27 | 35-37 | 38 | 39 | 40, 41 |
| Algeria | gallo_atl/gallo_med | 40, 41 | – | 42-50 | 51, 52 | 55-57 |
| North European<br>*M. edulis* | edu_eu/edu_am | 26-27 | – | 18, 19 | – | 10-17 |

For each hybrid zone, three analyses were carried out: (i) computation of hybrid indexes with the R package `introgress` (v1.2.3; Gompert and Buerkle, 2010), (ii) a local Bayesian clustering analysis using `Structure`, and (iii) a Bayesian genomic clines analysis.

`Structure` analyses for each hybrid zone used the same parameters as for the combined data set, and the admixture with linkage model. The filtration of markers was similar to the global analysis. Each subset was filtered for irrelevant genetic backgrounds in the hybrid zone studied. For example, in the Øresund hybrid zone we compare *M. edulis* (P1) and *M. trossulus* (P2). Therefore, *M. trossulus* individuals living in sympatry in the P1 populations were removed from this reference. Each hybrid zone was studied for a *K* of 2 and 3, to make sure there was no hidden substructure in the subset of individuals considered. The tros_am population was used as a distant reference for the Øresund hybrid zone, as the tros_eu_north population had a lower number of individuals and a higher level of missing data.

Genomic clines are used to detect markers deviating from the average genomic expectation given the distribution of hybrid indexes. The program `bgc` (v1.03) was used to estimate genomic cline parameters with a Bayesian method (Gompert & Buerkle, 2011, 2012) for each hybrid zone considered. Datasets were prepared with custom R scripts. For each hybrid zone, fixed markers were removed as they are uninformative. The same filtered marker sets as for the `Structure` analyses were used. Four independent chains of 200 000 iterations including 20 000 burn-in iterations with a thinning of 20 were performed. We used the ICARrho model for linked loci with the previously generated genetic map. The R package `rhdf5` (v2.32.0; Fischer, Pau, and Smith, 2019) was used to read the MCMC outputs. Convergence was assessed using the method and R code of Vehtari, Gelman, Simpson, Carpenter, and Bürkner (2019).

For each hybrid zone, we performed two analyses called “local” and “distant”. The local or distant parental populations are taken as references, respectively (Table 3). Both analyses considers the “central” population as admixed.

Loci exhibiting extreme deviations from the neutral genetic background were determined by two methods. First, we estimated locus-specific posterior distributions for the cline parameters *α* and *β*. Loci were classified as having “excess ancestry” if the 95% quantiles of these distributions did not include 0 (Gompert & Buerkle, 2012).

## 3 Results

We use a set of 212 differentiated and outlier markers to investigate the population genetics of several hybrid zones, and other introgressed populations in the *Mytilus* species complex. We start by showing that our target species and populations are identifiable with our ancestry-informative panel of SNPs. We also identify previously uncharacterised lineages or admixed clusters. Then we analyse the differentiation and introgression patterns in the hybrid zones, at two spatial scales.

### 3.1 Power of the SNP panel and clustering results

Our markers were taken from a previous GBS study (Fraïsse, Belkhir, et al., 2016), and so we first tested if the markers were still informative in our broader sample. To do this, we correlated *F*_*ST*_ values between the complete dataset (BeadXpress and GBS) and the GBS genotypes considered alone (Table S2). Results showed good delimitation (i.e. high *F*_*ST*_ correlations) between species and between known semi-isolated lineages within a species (e.g. each side of the Atlantic ocean or the Almeria-Oran front), while comparisons between less separated entities were less successful (i.e. low *F*_*ST*_ correlations). Overall, however, the assay design produced a strong enrichment in high *F*_*ST*_ markers compared to the distributions of Fraïsse, Belkhir, et al. (2016), showing that our markers could differentiate the lineages of interest (Figure S1).

Although the markers are informative, another consequence of increasing the sample size was that truly diagnostic markers became rare or absent at the species level. While our dataset contained many differentiated loci as expected, only two markers showed fixed differences between species. Indeed, when considering all reference individuals of one species, only the marker 147 was fixed between *M. edulis* and *M. galloprovincialis* (Contig H_L1_abyss_Contig244 position 6092 in Fraïsse, Belkhir, et al., 2016) and only the marker 015 was fixed between *M. edulis* and *M. trossulus* (Contig Contig17324_GA36A position 1089 in Fraïsse, Belkhir, et al., 2016).

Using our SNP panel, five genetic clusters could be defined without ambiguity in a Principal Component Analysis (Figures S3; and noting that the order of the PCs is not meaningful, due to our method of selecting SNPs). As shown in Figure 2, the three species are clearly differentiated, and so are known genetic clusters within each species: (i) European and American *M. edulis*, (ii) Atlantic and Mediterranean *M. galloprovincialis*, (iii) Baltic and American/North-European *M. trossulus*. We observed that the Baltic population of *M. trossulus* (22) is introgressed by *M. edulis*, as previously described (Fraïsse, Belkhir, et al., 2016), and this cluster can be identified with `Structure` with *K* = 8 (Figure S4). (Note that the placement of the clusters is not meaningful in Figure 2 due to the UMAP algorithm).

**Figure 2:**
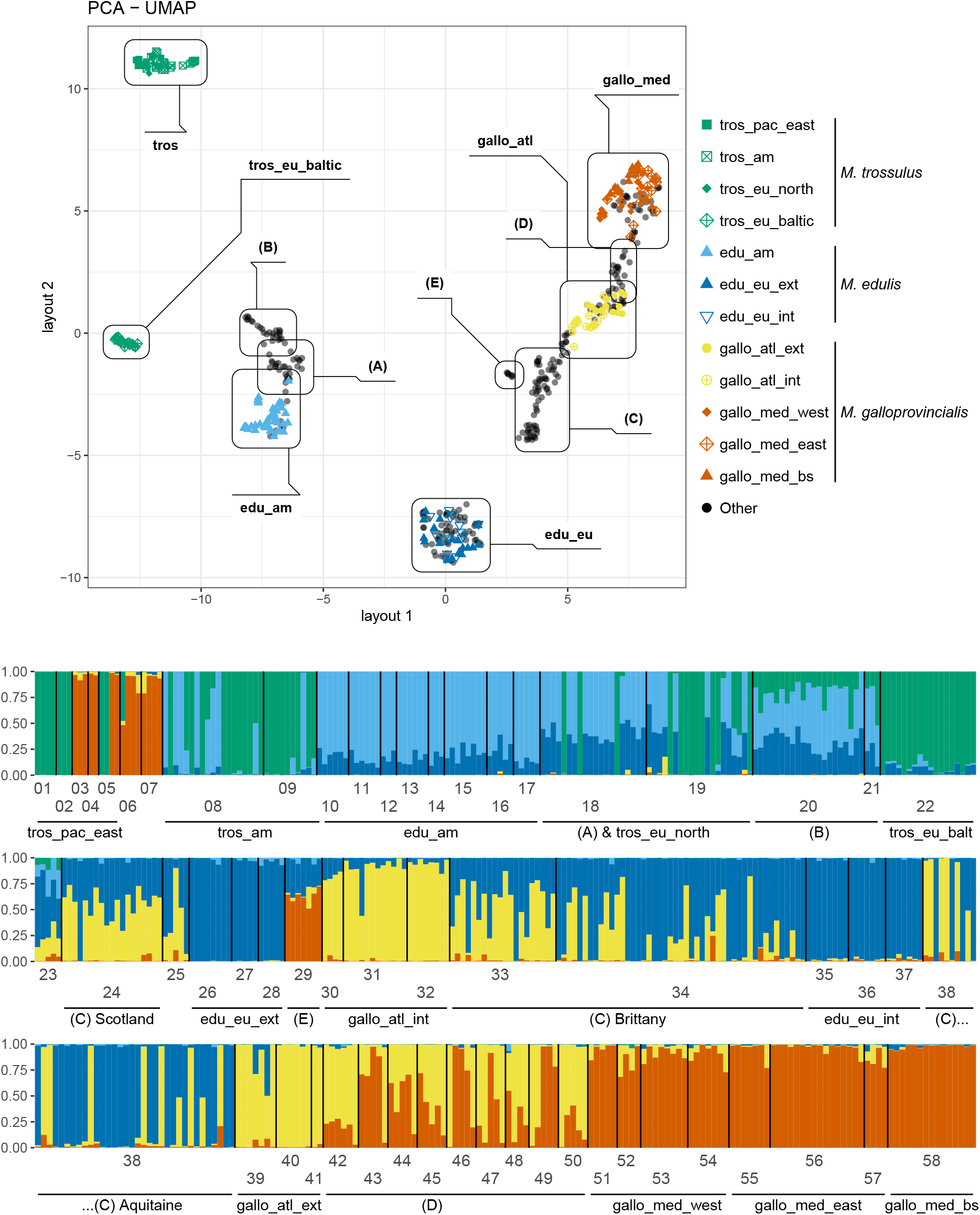
**top**: PCA-UMAP using the first 11 principal components. The reference level L3 (Table 1) is colour and shape coded. Note that this representation does not conserve distances and is designed to maximise groupings between similar entities, see Figure S3 for PCAs. Annotations show five groups of interest discussed in the main text: (A) North-European *M. edulis*; (B) Oresund hybrid zone; (C) Brittany, Aquitaine and Scotland hybrid zones (D) Algerian hybrid zone and (E) the port of Cherbourg. **bottom**: Ancestry composition of each individual in the dataset obtained with `Structure` for the major mode of *K* = 5. Populations are mainly ordered geographically. See Table S1 for population information.

The remaining individuals in Figure 2 form clouds labelled A to E and represent the various admixed populations we sampled (see also S3). For example, the hybrid zones of Brittany, Aquitaine and Scotland all involve Atlantic *M. galloprovincialis* and European *M. edulis*, and all are found in group C (Figure 2). By contrast, the Algeria hybrid zone involves Atlantic and Mediterranean *M. galloprovincialis*, and is group D (Figure 2).

Group E contains a small group of individuals sampled in the leisure marina of Cherbourg in 2003, on the French coast of the English Channel (29-Cherbourg). These individuals exhibited an admixture, largely between Mediterranean *M. galloprovincialis* (66% ancestry) and European *M. edulis*, (29% ancestry). We note that a nearby population, in Barfleur (population 28, around 30 km from Cherbourg), is composed exclusively of European *M. edulis*, the expected local genetic lineage.

Group A comprises North-European *M. edulis*. Those *M. edulis* individuals found in Russia (populations 18 and 19), in sympatry with *M. trossulus*, exhibit intermediate ancestries between South-European *M. edulis* (edu_eu_ext and int) and the American lineage of *M. edulis* (edu_am; Figures 2 and S10). Additionally, they do not differentiate on a secondary axis on a classic PCA (Figure S3). The ancestry intergradation observed in Figure S10 for populations 18 and 19 cannot be attributed to unaccounted *M. trossulus* ancestry, because those individuals do not exhibit introgression from this species in the global `Structure` analysis (Figure 2). A similar admixture is also visible in the Edinburgh population (23), which has additional Atlantic *M. galloprovincialis* ancestry due to its localisation close to the Scottish hybrid zone. Finally, group B corresponds to admixed individuals of the Øresund hybrid zone between the newly identified North-European *M. edulis* and the Baltic *M. trossulus* and populations 20-21 in Figure 2).

`Structure` analysis shows some further patterns. First, on the West coast of the USA (populations 01 to 07, Figure 2), we observe introduced Mediterranean *M. galloprovincialis* individuals, and a single F1 hybrid with the native *M. trossulus* parent (population 06). This is consistent with the report of Saarman and Pogson (2015).

Second, *M. edulis* samples from the Long Island region (USA, East coast, populations 10 to 16) were mainly assigned to the American *M. edulis* cluster (Figure 2), but there also appears to be some infra-specific ancestry coming from Europe (already noticed by Simon et al., 2020). An analysis considering only *M. edulis* samples, Figure S10, shows more clearly that individuals from the most southern populations of the Long Island Sound sometimes have higher European ancestry than other American populations (e.g. Boston, population 17).

### 3.2 Genetic differences between patches of parental populations close or distant to hybrid zones

We investigated the allele frequency differences between an intergradation of populations both between *M. edulis* and *M. galloprovincialis* (Figure 3), and between *M. edulis* and *M. trossulus* (Figure 4). This representation illustrates that outlier loci (identified by Fraïsse, Belkhir, et al., 2016) between populations within parental species most often correspond to a high frequency of the heterospecific allelic state (defined on the basis of allele frequency differences between species) in the population close to hybrid zones: blue smudges (edulis allelic state) in *M. galloprovincialis*, red smudges (galloprovincialis allelic state) in *M. edulis* (Figure 3), blue smudges (edulis allelic state) in *M. trossulus* and green smudges (trossulus allelic state) in *M. edulis* (Figure 4). The unbalance between populations close and distant to hybrid zones is indicative of differential introgression (similar reasoning to the now well established Patterson’s D or Reich’s f3), but at a given SNP the allelic state is not informative of the ancestry, and so this observation can also be explained by an increased frequency of a shared ancestral allele. However, (Fraïsse, Belkhir, et al., 2016) used gene phylogenies to identify that roughly half of these outliers were the consequence of enhanced local introgression while the other half was the consequence of a local selective sweep independent of introgression. Because the present study only uses strongly differentiated and outlier SNPs, we lack the neutral baseline required to re-conduct outlier tests done by Fraïsse, Belkhir, et al., 2016). Nevertheless, we confirm a strong signal that does not seem consistent with a fully neutral interpretation knowing the very low level of genetic differentiation between populations of the same species at the genome scale (Fraïsse, Belkhir, et al., 2016). We regret we cannot present this low differentiation baseline in the present paper and we can only refer the reader to the study of (Fraïsse, Belkhir, et al., 2016).

**Figure 3:**
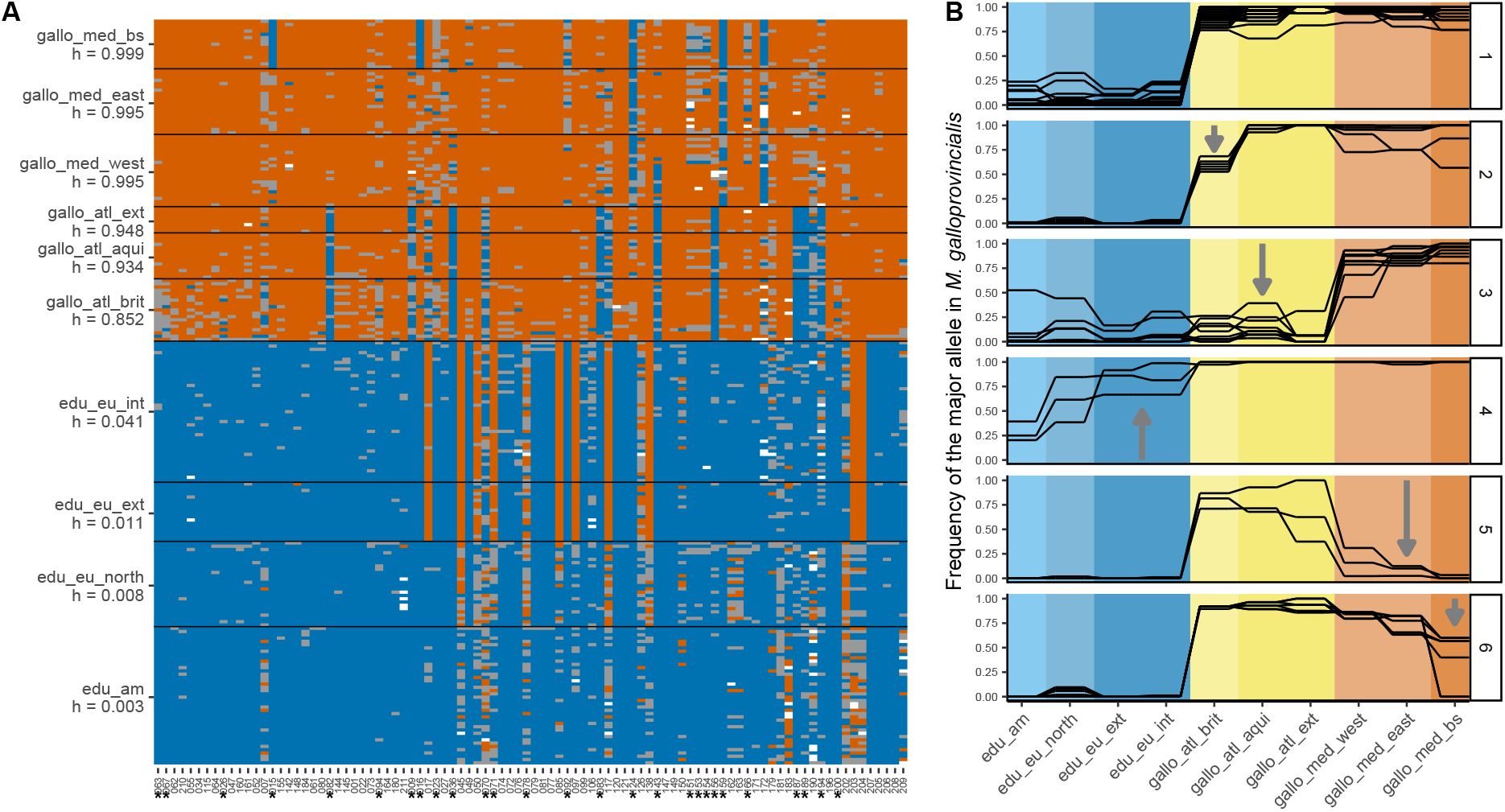
Visualisation of introgression between *M. galloprovincialis* and *M. edulis* using the reference dataset. **(A)** Plot of the raw genotypes. Orange: homozygous *M. galloprovincialis*; blue: homozygous *M. edulis*; grey: heterozygous. The orientation of alleles and mean hybrid indexes for each population (h) were computed with the R package `introgress`. The edu_eu_north population was taken as parent 1 (P1), and the gallo_med_east and gallo_med_bs populations were taken as parent 2 (P2). Loci identified by an asterisk are plotted in panels B2-6. **(B)** Allele frequency of the *M. galloprovincialis* allele (G) for a few loci selected visually to exemplify six categories found in the dataset: 1) loci informative between *M. edulis* and *M. galloprovincialis*; 2) introgression only in the local *M. galloprovincialis* Atlantic of Brittany; 3) introgression in all *M. galloprovincialis* Atlantic populations; 4) introgression in *M. edulis* European populations; 5) potentially old introgression only in the *M. galloprovincialis* Mediterranean; 6) potentially old introgression only in the *M. galloprovincialis* from the Black Sea.

**Figure 4:**
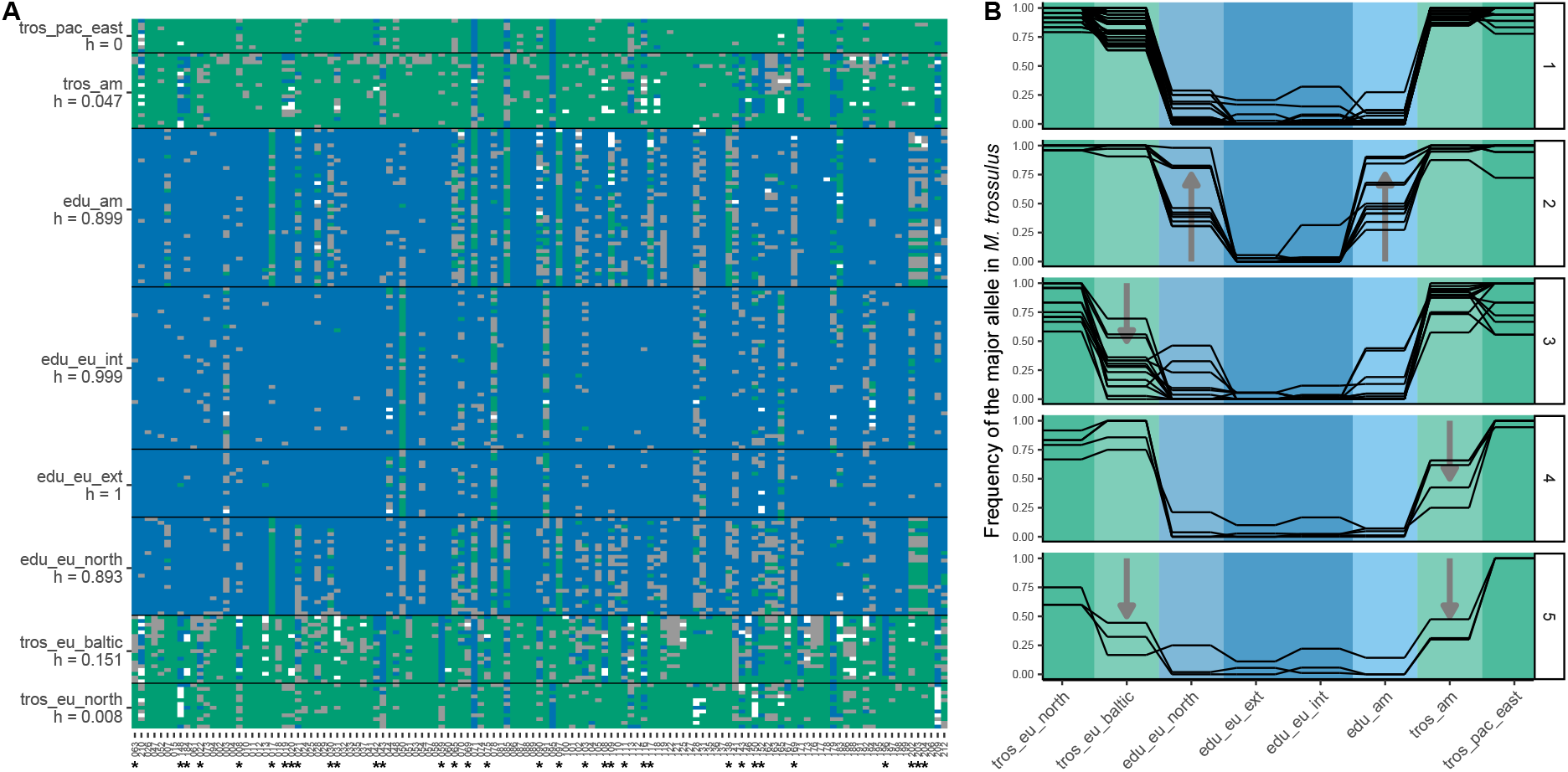
Visualisation of introgression between *M. trossulus* and *M. edulis* using the reference dataset. **(A)** Plot of the raw genotypes. Green: homozygous *M. trossulus*; blue: homozygous *M. edulis*; grey: heterozygous. The orientation of alleles and mean hybrid indexes for each population (h) were computed with the R package `introgress`. The tros_pac_east and tros_eu_north populations were taken as parent 1 (P1), and the edu_eu_ext and edu_eu_int ones were taken as parent 2 (P2). Loci identified by an asterisk are plotted in panels B2-5. **(B)** Allele frequency of the *M. trossulus* allele (T) for a few loci selected visually to exemplify five categories found in the dataset: 1) loci informative between *M. edulis* and *M. trossulus*; 2) introgression into *M. edulis* populations, either in both American and European or just one of them; 3) introgression into the Baltic *M. trossulus* population; 4) introgression into the East-American *M. trossulus* population; 5) introgression into East-American and Baltic *M. trossulus* populations, also showing a reduced introgression into the North-European *M. trossulus*.

The *M. edulis*-*M. galloprovincialis* mosaic hybrid zones (Figure 3) can be seen as a large scale intergradation between the Mediterranean populations (starting in the Black Sea) and the North-European ones. *M. galloprovincialis* populations are known to be more introgressed by *M. edulis* alleles in the Atlantic than the Mediterranean and our results confirm and illustrate this trend (see mean hybrid index, h, in Figure 3A). In addition, the local *M. galloprovincialis* population in Brittany (gallo_atl_brit), surrounded by two patches of *M. edulis* on either side, has a higher level of introgression than the local population in Aquitaine (gallo_atl_aqui). Interestingly, the Black-Sea population (gallo_med_bs) displays a few fixed alleles with an *M. edulis* allelic state, contrasting with the rest of the Mediterranean basin (Figure 3A and B6). The South-European *M. edulis*, both external and internal to the mosaic, have nearly as many smudges of *M. galloprovincialis* allelic state (Figure 3A and B4); and around half of these extends to Northern Europe.

For the *M. edulis*-*M. trossulus* comparison, we investigated two intergradations, in America and Europe (Figure 4). American and North European *M. edulis* populations show introgression from *M. trossulus*, with shared patterns of heterospecific alleles. Those two populations share potential local introgressions (Figure 4A), and almost none are private to either population (though see marker 183 in Figure 4). Regarding introgression from *M. edulis* to *M. trossulus*, whereas the Baltic *M. trossulus* has been highly introgressed (exhibiting a mean hybrid index of *h* = 0.15), American *M. trossulus* does show lower levels of global introgression (mean hybrid index of *h* = 0.05).

### 3.3 Concordance in hybrid zones

The hybrid zones studied here involve four pairs of lineages, as listed in Table 3. We considered three types of populations: (i) “central” populations, from the centre of the hybrid zone where two species or lineages coexist; (ii) local parental populations (P1 and P2 local) on each side of the hybrid zone and impacted by recent hybridisation; and (iii) distant parental populations (P1 and P2 distant) which are not in direct contact with the focal hybrid zone and so potentially less affected by local introgression.

A genomic cline analysis is designed to identify loci that shift from the genome wide expectation on the basis of the multi-locus hybrid index. Departure from the expectation is usually interpreted as evidence of selection (adaptive introgression, incompatibility or balancing selection). We posit that such analysis can strongly depend on the reference populations used in the analysis. Our outlier SNPs for instance should not appear as outlier in the genomic clines analysis if the local parental populations are used as references, but should if distant populations are used instead. We therefore contrast analyses done with different references.

As a general trend, the local genomic clines (Figure 5, middle panels) exhibited good convergence of the models in the Bayesian analyses (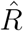 close to 1, Vehtari et al., 2019) with a limited number of excess ancestry markers both for *α* and *β*. On the other hand, analyses with distant parental populations taken as reference (Figure 5, right panels) exhibited an increase in outlier loci. We note that certain new outliers may be false positives due to a reduced convergence between chains in the later analysis.

**Figure 5:**
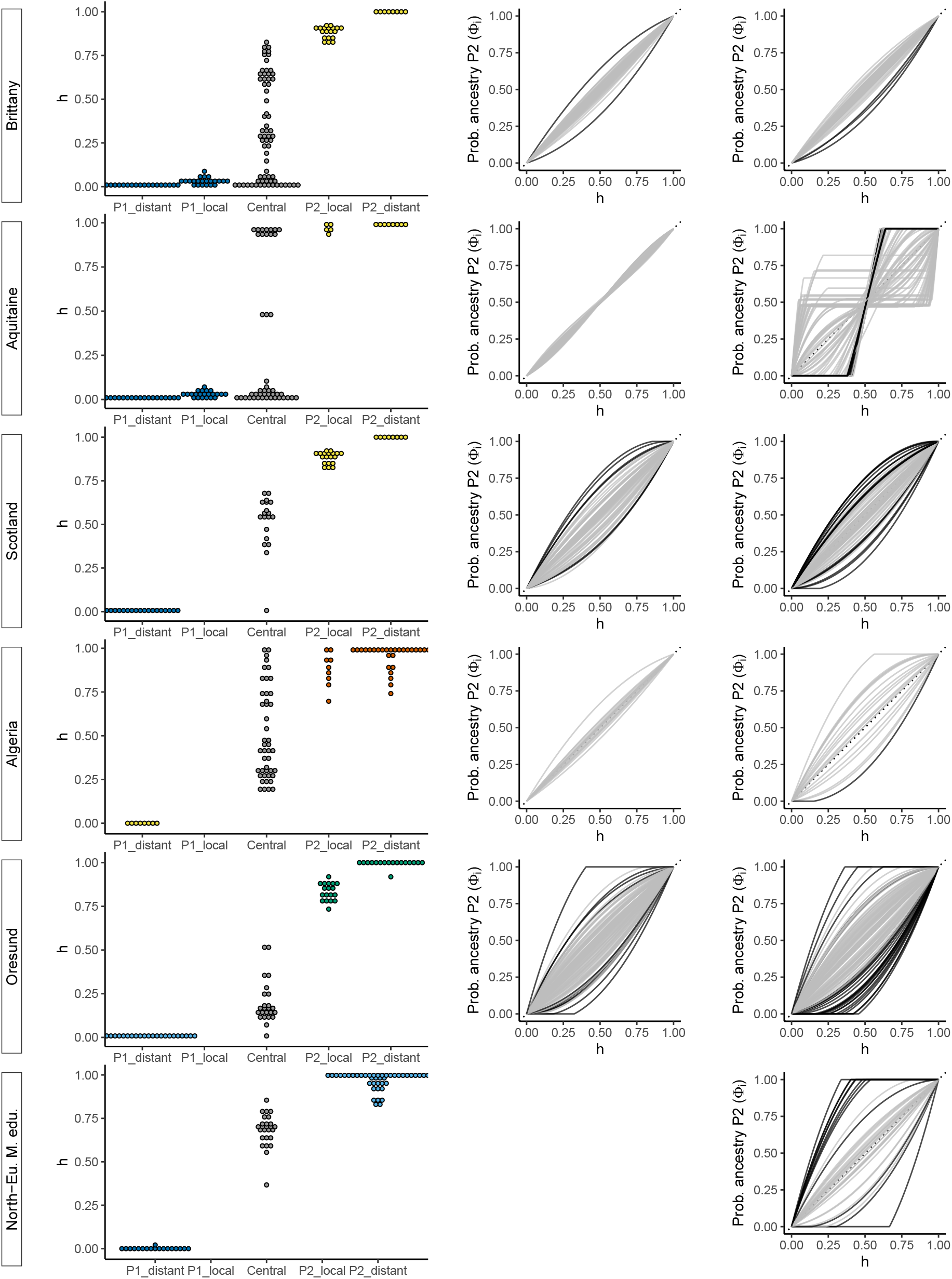
**Left panels:** Hybrid index (h) distributions from `introgress`, for each group of the hybrid zone. **middle panels:** Genomic clines computed with the local parental groups (P1_local, P2_local or distant when not available) with clines presenting an excess in either *α* or *β* parameters highlighted in black. **right panels:** Genomic clines computed with distant populations as parental groups. For genomic clines, only markers with an allele frequency difference > 0.3 between P1_periph and P2_periph are drawn. See Table 3 for a list of used populations.

Three hybrid zones, in Brittany, Aquitaine and Scotland, involve contact between South-European *M. edulis* and Atlantic *M. galloprovincialis*. The Brittany and Aquitaine zones are on one side of a patch of South-European *M. edulis* on the French Atlantic coast (Bierne, Borsa, et al., 2003). Nevertheless, these two zones exhibit strong differences in their hybrid index distributions, and this is reflected in their genomic clines (Figure 5). On one hand, Brittany presents a broad distribution of hybrid types between the two parental populations and a local *M. galloprovincialis* parental population that is introgressed (P2_local, see also Figure S6). On the other hand, Aquitaine presents only three F1 hybrids between the two parental types and the local *M. galloprovincialis* population (P2_local, including some individuals in sympatry with the *M. edulis* P1_local population) does not present strong introgression compared with the distant population (P2_distant). In Scotland, the central population exhibits variable hybrid indexes, and so resembles Brittany more than Aquitaine. The genetic composition of local *M. edulis* and *M. galloprovincialis* populations in Scotland are largely unknown. However the only *M. edulis* individual sampled in population 24 is indiscernible from the edu_eu_ext references (Figures 2 and 5).

The Algerian hybrid zone was sampled most extensively (El Ayari et al., 2019). As previously shown with 4 markers (El Ayari et al., 2019), the distribution of hybrid indexes is wide, and the zone of coexistence of the two lineages extends to around 600 km. We did not sample Atlantic *M. galloprovincialis* populations close to this zone (P1_local). If the local population and hybrid zone were harbouring local introgressions, we would have expected discordant loci when using the P1_distant population as parental. As this is not the case, we can hypothesise that the local population of Atlantic *M. galloprovincialis* is not heterogeneously introgressed by the Mediterranean *M. galloprovincialis*.

The Øresund hybrid zone includes the North-European *M. edulis* lineage (Figure 2, populations 20-21). Therefore, we treated such individuals from Russia as the P1 parental population. In this zone, the central population exhibits relatively homogeneous hybrid indexes, and is mainly composed of North-European *M. edulis* introgressed by *M. trossulus* (Figure 2). On the other side of the hybrid zone, the Baltic *M. trossulus* population is introgressed as shown above (Fraïsse, Belkhir, et al., 2016; Väinölä and Strelkov, 2011; Figures 4 and 5).

To highlight the intermediate character of the North-European *M. edulis* lineage, we carried out an analysis treating as admixed the North-European *M. edulis* individuals in the Russian populations (18 and 19, Fig. S10). As shown by the hybrid index and the `Structure` analyses, these mussels have an homogeneous admixture of around 60% South-European *M. edulis* and 40% American *M. edulis*.

When finally considering the correlation of genomic cline parameters (*α* and *β*) between hybrid zones, only two correlations proved statistically significant after correcting for multiple tests. The correlated parameters are *α* and *β* between the Scotland and Brittany local hybrid zone (Spearman correlation coefficients of 0.38 and 0.36, respectively, with *p*-values < 0.005), and *β* between the Brittany and Aquitaine hybrid zones (Spearman correlation coefficient of 0.38, *p*-value = 0.002).

## 4 Discussion

### 4.1 *Mytilus* mussels are genetically differentiated, but fixed differences are extremely rare

One aim of this study was to develop a panel of highly differentiated, and potentially diagnostic markers for the *Mytilus* complex in the Northern Hemisphere. To this end, we started with 51,878 high-quality SNPs from 1269 sequences (Fraïsse, Belkhir, et al., 2016), and selected SNPs with the greatest discriminatory power. We combined differentiated variants between species with intra-specific *F*_*ST*_ outliers to provide a more detailed identification. This procedure was successful, in that we were able to discriminate not only individuals of the three species (*M. edulis*, *M. galloprovincialis* and *M. trossulus*) but also partially isolated genetic lineages within species (European and American *M. edulis*; Atlantic and Mediterranean *M. galloprovincialis*; and Baltic and American/North-European *M. trossulus*; Figure 2).

Nevertheless, of 92 highly differentiated loci between *M. edulis* and *M. galloprovincialis* (Figure 3), only one was a fixed difference in our sample. Similarly, there was only one fixed difference among 125 highly differentiated loci between *M. edulis* and *M. trossulus* (Figure 4). All the other SNPs were found at least once in heterozygous state (grey squares in Figures 3 and 4) and even sometimes as heterospecific homozygotes. Our approach highlights that loci originally claimed to be diagnostic are likely to loose their diagnostic status in a large-scale assay.

Similar failures of diagnostic loci in *Mytilus* have already arisen for the widely used Me15/16 marker (also known as Glu-5’) where the “E” allele was observed in *M. galloprovincialis* (Bierne, Borsa, et al., 2003; Borsa, Daguin, Ramos Caetano, & Bonhomme, 1999; Hamer, Korlević, Durmiši, Nerlović, Bierne, et al., 2012; Wood, Beaumont, Skibinski, & Turner, 2003) and the “G” allele in *M. edulis* (Bierne, Borsa, et al., 2003; Kijewski, Wijsman, Hummel, & Wenne, 2009; Luttikhuizen, Koolhaas, Bol, & Piersma, 2002). In *Mytilus* mussels, therefore, it seems prudent to replace single marker diagnosis with multilocus inference; and our results suggest that 5-10 well chosen loci should be sufficient for this purpose, but not less.

It is unlikely that this situation is unique to mussels. In the highly divergent *Ciona* tunicate species, *C. robusta* and *C. intestinalis*, loci initially assumed to be diagnostic and used to identify hybrids (Bouchemousse, Lévêque, Dubois, & Viard, 2016; Nydam & Harrison, 2011; Sato, Shimeld, & Bishop, 2014) were subsequently found to harbour shared polymorphisms in a multilocus analysis, and heterozygous genotypes were found in parental populations, both at the initially studied markers and at many of the newly designed markers (Bouchemousse, Liautard-Haag, Bierne, & Viard, 2016). As for the *Mytilus* system, *Ciona* species have very high nucleotide diversities and have been shown to go through a history of secondary contact (Roux, Tsagkogeorga, Bierne, & Galtier, 2013) resulting in a higher rate of shared and high frequency private polymorphisms than expected at first sight for well isolated and low *N*_*e*_ species.

### 4.2 Enhanced local introgression is widespread and has several possible causes

A notable feature of our data was the appearance of heterospecific alleles that were present at high frequency in one or a small number of populations. These alleles were very common (at least one such allele was found in each population of each lineage of all three species), but were found heterogeneously across the genome (Figures 3 and 4).

Fraïsse, Belkhir, et al. (2016) also found this pattern, in their much smaller sample. Furthermore, their use of genome-wide data, combined with analyses of gene-genealogies, suggested that the pattern was due to local introgression, rather than, say, incomplete lineage sorting (e.g. the local sweep of a shared ancestral variant). While our data do not allow us to distinguish between these possibilities, and while both have no doubt taken place, our use of the data of Fraïsse, Belkhir, et al. (2016) to choose our SNPs, combined with the finding that these alleles are preferentially found in contiguous populations (Figures 3 and 4), suggest strongly that there are high levels of local introgression in the *Mytilus* complex (Fraïsse, Haguenauer, et al., 2018; Fraïsse, Roux, Gagnaire, et al., 2018).

The biogeography of the complex also allowed us to study introgression of a very particular kind. More specifically, the *M. galloprovincialis*/*M. edulis* transition is characterised by near-continuous intergradation between the Black Sea and Scandinavia, but with genetic barriers to gene flow at multiple points. This leads to a mosaic distribution in several regions. For example, the isolated *M. edulis* and *M. galloprovincialis* patches on the Atlantic coast of France, are separated by three hybrid zones, in Aquitaine, South Brittany, and Normandy (not sampled in this study) (Bierne, Borsa, et al., 2003; Hilbish et al., 2012). This structure revealed multiple instances of enhanced local introgression, that is, introgression events that are localised both genomically, and geographically, with allelic variants crossing one barrier to gene flow, but halted at a subsequent barrier (Figure 3).

Three possible mechanisms could explain the heterogeneity of local introgression. First, introgression could be adaptive (Hedrick, 2013; Pardo-Diaz et al., 2012; Racimo, Marnetto, & Huerta-Sánchez, 2017; Staubach et al., 2012). In this case, the fact that introgression was halted at a subsequent barrier could be explained either by an environmental difference (such that the allele was only locally beneficial), or by a very strong barrier (in which case the halting would be transient). However, Fraïsse, Belkhir, et al. (2016) found no clear links between the biological function of outlier genes and the environmental conditions of introgressed populations. Alternatively, if the markers are not the direct target of selection – as seems highly plausible (Fraïsse, Belkhir, et al., 2016) – a marker might hitchhike through one barrier, but be halted at a second (Barton, 2000; Faure et al., 2008).

A second possibility is that introgression acts to reduce genetic load in the recipient population (Harris & Nielsen, 2015; Juric, Aeschbacher, & Coop, 2016; Kim, Huber, & Lohmueller, 2018). This scenario is similar to adaptive introgression, but emphasises the role of deleterious mutations in the recipient population rather than advantageous mutations in the donor population.

Third, and finally, introgression could involve bi-stable variants in a tension zone (Barton & Hewitt, 1985). Such variants can individually move via an asymmetry in parental fitness, i.e. one parental genotype is fitter than the other, but both are fitter than hybrids (Barton, 1979a; Barton & Turelli, 2011). However they can easily be trapped for long periods by a density trough, a barrier to dispersion, and coupling (Barton and Turelli, 2011; Bierne, Welch, Loire, Bonhomme, and David, 2011, see El Ayari et al., 2019 for a possible example in *Mytilus*). This scenario also implies transience in the structure, because tension zones can move due to genetic drift or changes in the environmental conditions (Piálek & Barton, 1997). In other words, rare stochastic events can free the “pushed” wave of advance (Piálek & Barton, 1997). This hypothesis was already proposed in *Mytilus* by Fraïsse, Roux, Welch, and Bierne (2014, July), and has also been proposed for other systems such as in *Bathymodiolus* mussels (Plouviez et al., 2013), Sea Bass (Duranton et al., 2020) and Killer Whales (Foote et al., 2019).

### 4.3 High overall concordance of barrier loci at hybrid zones

With high levels of local introgression, it can be difficult to study genomic clines. With our data this was evident in the occasional failures of our cline analyses. Nevertheless, in most cases these analyses yielded clear results: a high level of concordance with respect to local parental populations (Figure 5 middle panels), and increased discordance with respect to distant parental populations (again, consistent with local introgression).

Even with local parental populations, we observed a few SNPs that deviated from the genomic average. The anomalous “excess ancestry” loci observed in Scotland and Øresund in the local analysis could still be a consequence of enhanced local introgression, as P1_local populations were missing from our sampling. Nonetheless deviating loci are still observed, e.g. in Brittany (Figure 5). These loci tend to introgress more than the others in the centre of the zone, but – unlike established introgression – have not escaped the genetic barrier. We suspect that these loci might contribute to bi-stable variation that is asymmetric, as described above, but with a lower fitness difference between parental genotypes.

Genomic cline outliers with asymmetric introgression are often interpreted as evidence of adaptive introgression, i.e. directional selection (Gompert & Buerkle, 2009). Our results support directional selection in the centre of the hybrid zones, but not in flanking parental populations. If a force is opposing the spread of the adaptive variant from a barrier, it could be selection against heterozygous or recombinant hybrid genotypes (Barton, 1979b).

Finally, the observation of a small but sometimes significant correlation of the genomic cline parameters can point us to parallel processes that may be at play in the hybridisation of *Mytilus* species. Here, correlations where only found in hybrid zones implicating the same species and lineages. However, Simon et al. (2020) already showed that parallel deviations in admixed populations of mussels exist, and that it can also sometime happen between different lineages. Such cases call for further investigations of potential selective effects in the regions of the studied markers, either due to positive selection or barrier loci.

### 4.4 The timing and context of introgressions

Given the heterogeneity in the introgression patterns we have observed, it seems likely that they were shaped by both contemporary contacts, and historic contacts during the Quaternary period (Hewitt, 2000).

In some cases our data allow us to make inferences about the timing of specific introgression events. For example, local introgression of *M. edulis* alleles into *M. galloprovincialis* from the Black Sea and Mediterranean, is consistent with ancient contacts between these populations (Fraïsse, Belkhir, et al., 2016; Gosset & Bierne, 2013). The two private introgressions into the Black sea were nevertheless unexpected. Fraïsse, Belkhir, et al. (2016) did not sample these populations, and so our data set was not enriched for discriminatory SNPs. Therefore such introgressions might be quite numerous in the rest of the genome.

By contrast with these putatively ancient introgressions, it is likely that the private introgressions found in *M. galloprovincialis* from Brittany (Figure 3B2) are relatively recent. This is because, given its position enclosed in the mosaic, it is probable that this population only became established after the last post-glacial period.

More recently still, it is likely that the admixed individuals between Mediterranean *M. galloprovincialis* and South-European *M. edulis*, observed in the port of Cherbourg, represent a recent human-mediated introduction. Similar admixed individuals have been observed in other ports in the English Channel and French Atlantic coast (Simon et al., 2020), where these “dock mussels” form small-scale hybrid zones with the native lineages outside of the port.

In one final case, shared introgressions allow us to make inferences about the historical biogeography of the *Mytilus* complex. In particular, we found that several introgressions from *M. trossulus* to *M. edulis* were shared between *M. edulis* populations in Northern-Europe and America, and the most parsimonious explanation is that these introgressions predated their split. This hypothesis sheds light on the origins of the North-European *M. edulis* population. We have shown that this population is differentiated from both American, and South-European *M. edulis*, and appears as intermediate in the PCA (Figure S3), `Structure` (Figures 2 and S10) and hybrid index analyses (Figure 5). This observation previously made in Simon et al. (2020), has recently been confirmed by Wenne et al. (2020). Given its presence as the parental *M. edulis* population in the Øresund hybrid zone, and the complete absence of American ancestry in the Netherlands, the border between Southern and Northern European *M. edulis* probably falls somewhere near to the Danish coast (see also the previously observed differences between North Sea and populations North of the Kattegat region (Bierne, Daguin, Bonhomme, David, & Borsa, 2003; Stuckas, Stoof, Quesada, & Tiedemann, 2009)). Given the shared introgressions, therefore, the contemporary biogeography could reflect a recolonisation of America after the last glacial maxima, by a proto-North-European *M. edulis*, which was later introgressed by the South-European *M. edulis* (Wares & Cunningham, 2001). Alternatively, American *M. edulis*, having survived in a refugia (Riginos & Henzler, 2008), might have colonised the North-Atlantic and Scandinavia, and then been introgressed by the South-European *M. edulis* in Europe.

### 4.5 Implications for speciation research

The *Mytilus* species complex is characterised by diverse levels of genetic differentiation between lineages and species, and by pervasive introgression due to a dense history of secondary contacts and mosaic geographical distribution. As such, it constitutes a powerful model to investigate the maintenance of species barriers in the marine environment. Based on our results and previous studies (Bierne, Borsa, et al., 2003; Fraïsse, Belkhir, et al., 2016), we suggest that secondary contacts are not only periods of neutral introgression; but also periods during which genetic incompatibilities can be swamped resulting in a punctuated decay of the barrier to gene flow. This vision is rooted in the observation that gene flow after secondary contacts in the *Mytilus* complex is shown to be heterogeneous along the genome (Fraïsse, Roux, Gagnaire, et al., 2018; Roux et al., 2014; Roux et al., 2016). In regions of higher gene flow, some alleles have locally introgressed. Those can either be missed when distant parental populations are not used, or observed and interpreted as local adaptive introgression. Another explanation exists, considering that bi-stable variants, which could be associated with or responsible for genetic incompatibilities, can punctually escape a zone of coupling. Overall our results suggest that asymmetrical parental fitness differences may enhance introgression at some regions of the genome yet successive barriers can prevent or delay propagation.

## Data availability

All data and analyses pipelines will be deposited in an open archive upon acceptance (most probably Zenodo).

## Acknowledgements

Data used in this work were partly produced through the genotyping and sequencing facilities of ISEM and LabEx CeMEB, an ANR “Investissements d’avenir” program (ANR-10-LABX-04-01) This project benefited from the Montpellier Bioinformatics Biodiversity platform supported by the LabEx CeMEB. We thank Norah Saarman, Grant Pogson, Célia Gosset and Pierre-Alexandre Gagnaire for providing samples. This work was funded by a Languedoc‐Roussillon “Chercheur(se)s d’Avenir” grant (Connect7 project). P. Strelkov was supported by the Russian Science Foundation project 19-74-20024. This is article 2020‐XXX of Institut des Sciences de l’Evolution de Montpellier.

## Supplementary information

**Table S1:** Sampling information per population. pop: population name in the dataset; N: number of individuals genotyped from this population; lat: latitude (in decimal degrees); long: longitude (in decimal degrees); locality: details on the locality.

| pop | N | lat | long | locality |
| --- | --- | --- | --- | --- |
| 01-California-NEW | 4 | 33.609786 | -117.92158 | Newport Harbor, California, USA |
| 02-Oregon-Coos | 3 | 43.366 | -124.217 | Coos Bay, Oregon, USA |
| 03-California-Carlsbad | 3 | 33.156302 | -117.352779 | Carlsbad, California, USA |
| 04-SantaBarbara | 2 | 34.42 | -119.698 | Santa Barbara, California, USA |
| 05-California-TBR | 4 | 37.872228 | -122.451734 | Tiburon Point, SF, California, USA |
| 06-California-BRK | 4 | 37.859163 | -122.308652 | Berkeley Marina, SF, California, USA |
| 07-California-LAM | 4 | 37.802713 | -122.257995 | Lake Merritt, SF, California, USA |
| 08-StLawrence-CBD | 19 | 48.269414 | -69.466146 | Cap de Bon Désir, Québec, Canada |
| 09-StLawrence-TAD | 10 | 48.134413 | -69.696141 | Tadoussac, Québec, Canada |
| 10-Lgls-SHM | 6 | 41.386917 | -71.519167 | Snug Harbor Marina, Charlestown, Long Island, USA |
| 11-Lgls-QNT | 6 | 41.332694 | -71.713333 | Quonochontaug, Long Island, USA |
| 12-Lgls-BYY | 3 | 41.346694 | -71.967583 | Brewer Yacht Yard, Mystic River, Long Island, USA |
| 13-Lgls-OST | 6 | 41.263278 | -72.384194 | Old Saybrook Town, Hartlands Drive, Long Island, USA |
| 14-Lgls-BFM | 3 | 41.260556 | -72.820722 | Branford Harbor, Long Island, USA |
| 15-Lgls-MYC | 8 | 41.210917 | -73.051528 | Milford Yacht Club, Long Island, USA |
| 16-Lgls-SPM | 5 | 41.107222 | -73.354778 | Southport Marina, Long Island, USA |
| 17-Boston | 5 | 42.36 | -71.05 | Boston, USA |
| 18-NovyMost | 20 | 68.904723 | 33.026324 | Novy Most, Russia |
| 19-Retinskoye | 20 | 69.114088 | 33.387851 | Retinskoye, Russia |
| 20-Oresund-Helsinborg | 21 | 56.04 | 12.694 | Helsinborg, Oresund, Sweden |
| 21-Oresund-Raa | 3 | 55.999 | 12.74 | Raa, Oresund, Sweden |
| 22-Tvarminne | 18 | 59.841 | 23.201 | Tvarminne, Finland |
| 23-Edinburgh | 5 | 55.95 | -3.18 | Edinburgh, Scotland, UK |
| 24-Aberdeen | 19 | 57.14 | -2.094 | Aberdeen, Scotland, UK |
| 25-Kenmare | 5 | 51.88 | -9.583 | Kenmare, Ireland |
| 26-Wadden-sea | 9 | 53.31 | 5.424 | Wadden Sea, The Netherlands |
| 27-Calais | 5 | 50.974978 | 1.870983 | Calais, France |
| 28-Barfleur | 5 | 49.670616 | -1.263 | Barfleur, France |
| 29-Cherbourg | 7 | 49.63942 | -1.61994 | Cherbourg, France |
| 30-Dinard | 4 | 48.633 | -2.05 | Dinard, France |
| 31-Roscoff | 12 | 48.72 | -3.98 | Roscoff, France |
| 32-Guillec | 8 | 48.687003 | -4.071535 | Guillec, France |
| 33-Kerbihan | 20 | 47.576883 | -3.021033 | Kerbihan, France |
| 34-Pornichet | 47 | 47.26 | -2.34 | Pornichet, France |
| 35-Gascogne | 8 | 46.325 | -1.312 | Gulf of Gascogne, France |
| 36-Biscay | 7 | 45.95 | -1.032 | Bay of Biscay, France |

Table S1 – Continued from previous page
| pop | N | lat | long | locality |
| --- | --- | --- | --- | --- |
| 37-Aiguillon | 7 | 44.65 | -1.13 | Aiguillon, France |
| 38-Arcachon | 44 | 44.589 | -1.239 | Arcachon, France |
| 39-Biarritz | 7 | 43.483 | -1.558 | Biarritz, France |
| 40-Faro | 7 | 37.01 | -7.93 | Faro, Portugal |
| 41-Dahkla | 2 | 23.72 | -15.93 | Dahkla, Morocco |
| 42-Nador | 6 | 35.166 | -2.93 | Nador, Morocco |
| 43-Arzew | 5 | 35.85 | -0.316 | Arzew, Algeria |
| 44-Moustaganem | 5 | 35.93 | 0.083 | Moustaganem, Algeria |
| 45-SidiLakhdar | 5 | 36.165 | 0.439 | Sidi Lakhdar, Algeria |
| 46-ElMarsa | 5 | 36.81 | 3.244 | El Marsa, Algeria |
| 47-Ghouraya | 5 | 36.56 | 1.899 | Ghouraya, Algeria |
| 48-Tipaza | 5 | 36.49 | 2.381 | Tipaza, Algeria |
| 49-SidiFredj | 5 | 36.75 | 2.846 | Sidi Fredj, Algeria |
| 50-Bejaia | 5 | 36.75 | 5.0567 | Bejaia, Algeria |
| 51-Ziama | 5 | 36.66 | 5.483 | Ziama, Algeria |
| 52-Collo | 5 | 37 | 6.56 | Collo, Algeria |
| 53-Thau | 8 | 43.4 | 3.7 | Thau, France |
| 54-Nules | 7 | 39.853 | -0.155 | Nules, Spain |
| 55-Bizerte | 7 | 37.26 | 9.86 | Bizerte, Tunisia |
| 56-Heraklion | 16 | 35.338 | 25.1442 | Heraklion, Greece |
| 57-Croatia | 4 | 44.43 | 14.979 | Croatia |
| 58-Trabzon | 15 | 41 | 39.71 | Trabzon, Turkey |

**Table S2:** Linear models of *F*_*ST*_ comparisons between genotype-by-sequencing (GBS) and both methods combined. For each comparison, the Weir and Cockerham (1984) *F*_*ST*_ is computed for each locus and then correlated between the full dataset and the GBS only dataset.

| Level | Comparison | slope (p-value) | intercept (p-value) | adjusted $r^2$ | p-value model |
| --- | --- | --- | --- | --- | --- |
| L1 | edu/gallo | 0.93** | 2.74e-2* (4.28e-3) | 0.96 | ** |
|  | edu/tro | 0.95** | 2.07e-2 (7.47e-2) | 0.95 | ** |
|  | gallo/tro | 0.93** | 3.40e-2* (1.72e-2) | 0.92 | ** |
| L2 | tro am/eu | 0.48** | 4.74e-2* (5.04e-4) | 0.66 | ** |
|  | edu am/eu | 0.85** | 1.27e-2 (0.33) | 0.85 | ** |
|  | gallo atl/med | 0.90** | 1.04e-3* (7.2e-06) | 0.87 | ** |
| L3 | edu eu ext/int | 0.46** | 1.54e-2 (9.73e-2) | 0.59 | ** |
|  | gallo atl ext/int | 0.46** | 2.65e-2 (4.84e-3) | 0.45 | ** |
|  | gallo med east/west | 0.23** | 1.46e-2* (2.28e-2) | 0.47 | ** |
|  | edu intra am | 0.39** | -5.94e-3 (0.45) | 0.71 | ** |

\*\* : significant with $p$ -value $< 2.2e-16$ ; \* : significant with indicated $p$ -value.

**Figure S1:**
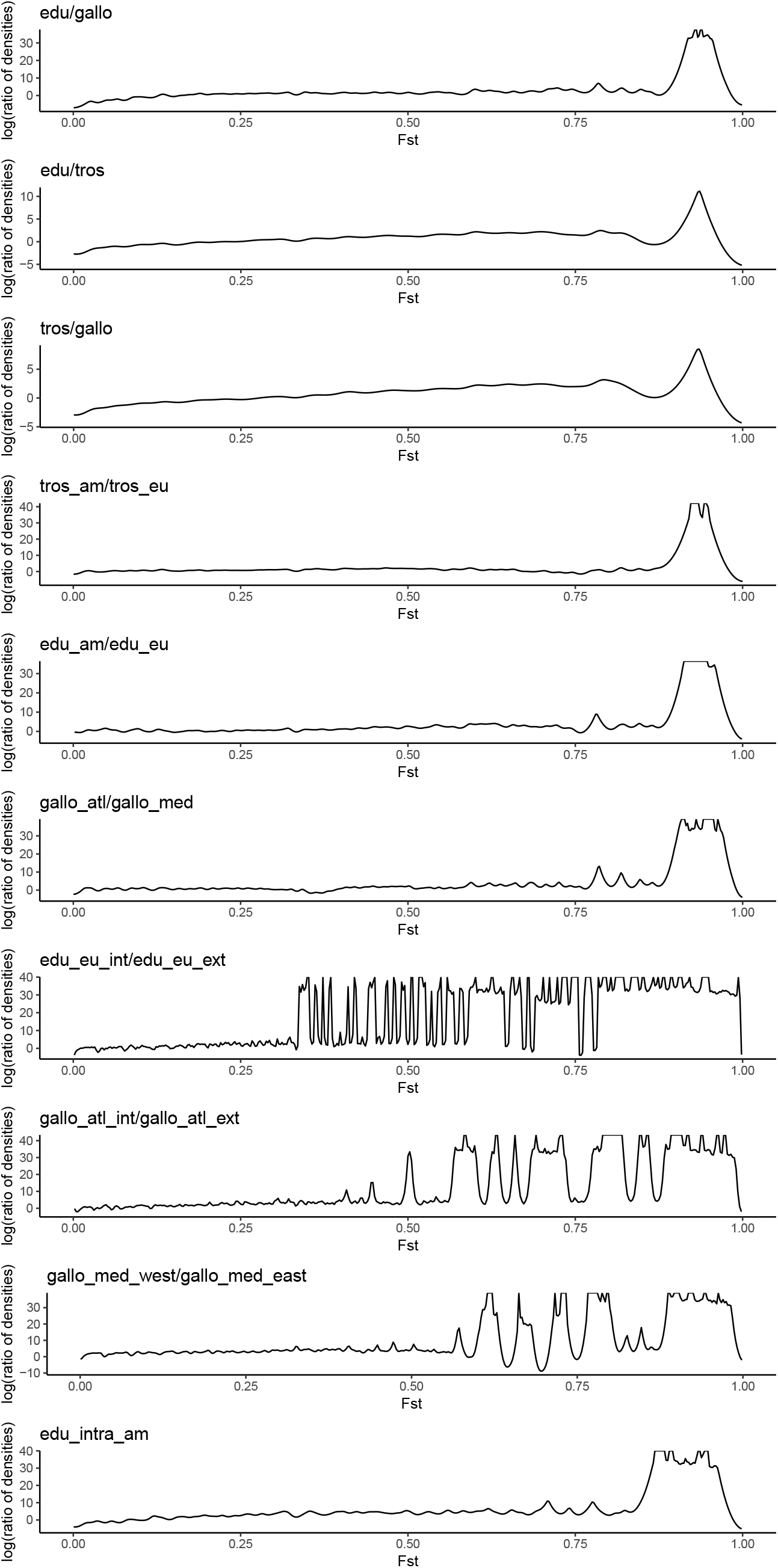
Enrichment in high *F*_*ST*_ markers for all comparisons. y-axis: natural logarithm of the ratio of *F*_*ST*_ densities between the ancestry informative dataset (present study) and the complete dataset (Fraïsse, Belkhir, Welch, & Bierne, 2016). Densities for *F*_*ST*_ between 0 and 1 are presented, so that negative *F*_*ST*_ values are omitted.

**Figure S2:**
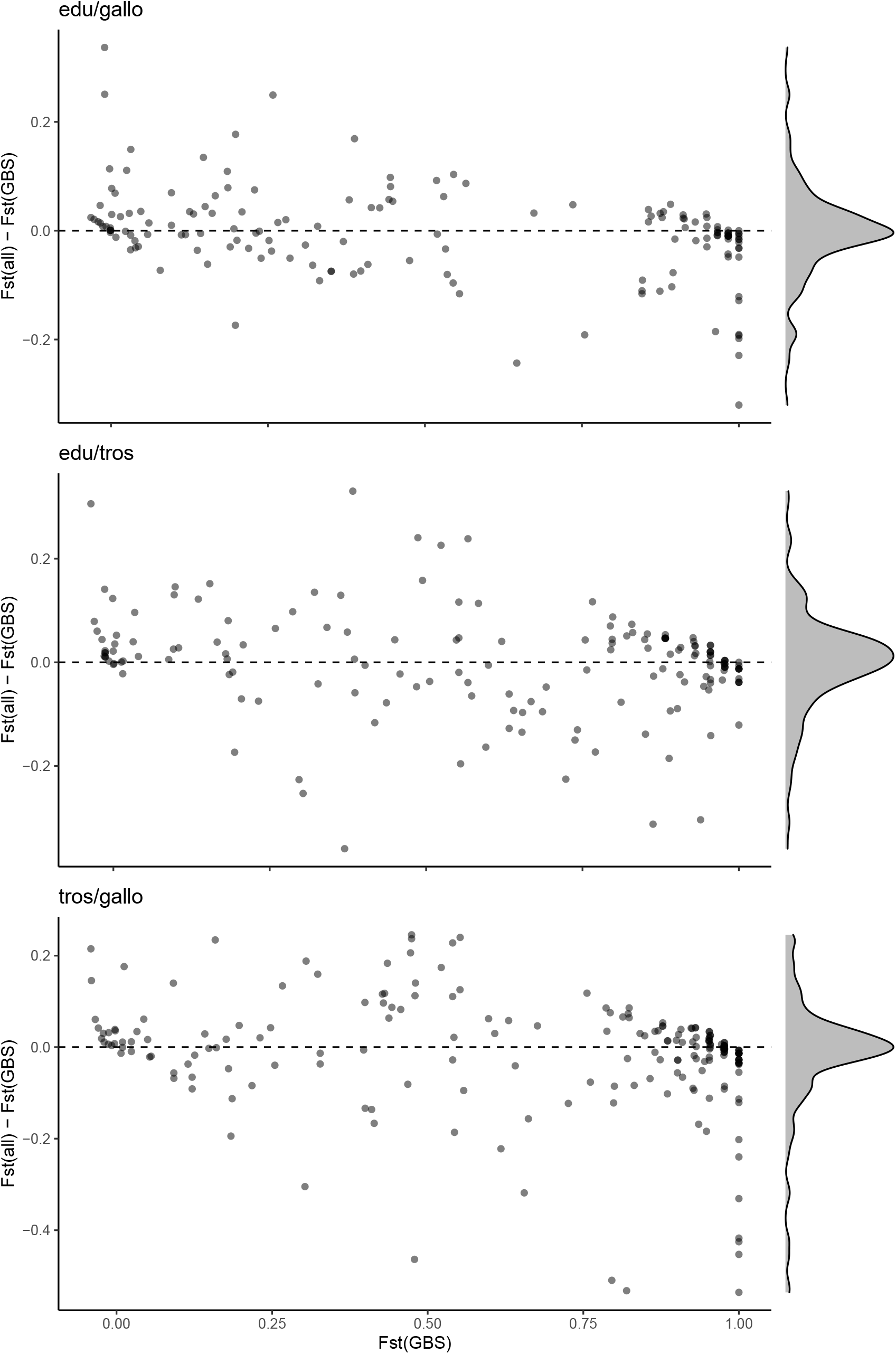
*F*_*ST*_ difference between the complete dataset and the GBS only dataset plotted against the *F*_*ST*_ of the GBS dataset, for each comparison of species. This figure shows the variance in *F*_*ST*_ produced by the increase of sampling.

**Figure S3:**
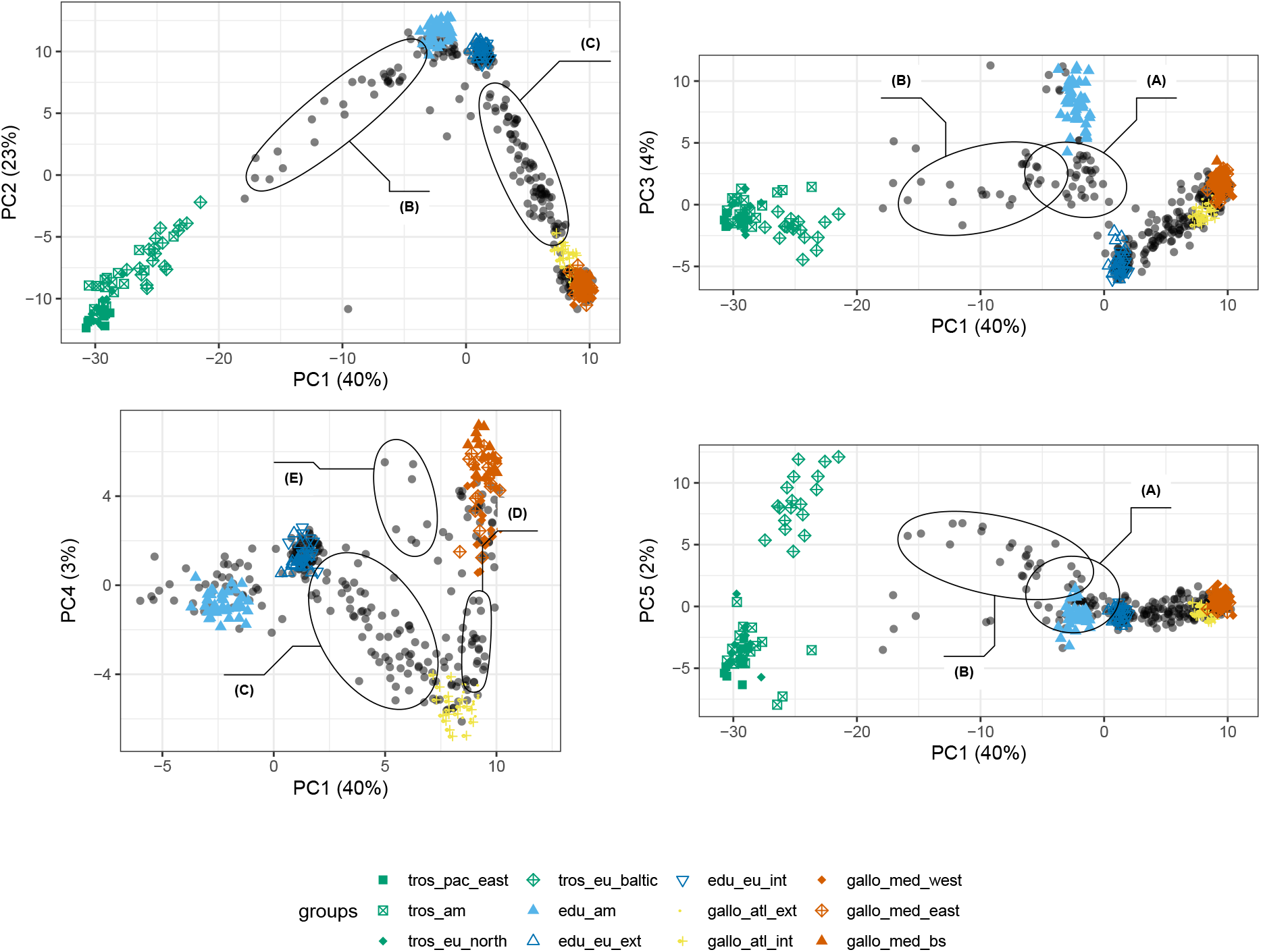
Principal Components Analysis presented for the first 5 components. In the bottom left PCA, *M. trossulus* individuals were cut out from the plot to zoom on the *M. edulis* and *M. galloprovincialis* interesting parts. Annotations show five groups of interest discussed in the main text: (A) North-European *M. edulis*; (B) Oresund hybrid zone; (C) Brittany, Aquitaine and Scotland hybrid zones (D) Algerian hybrid zone and (E) the port of Cherbourg.

**Figure S4:**
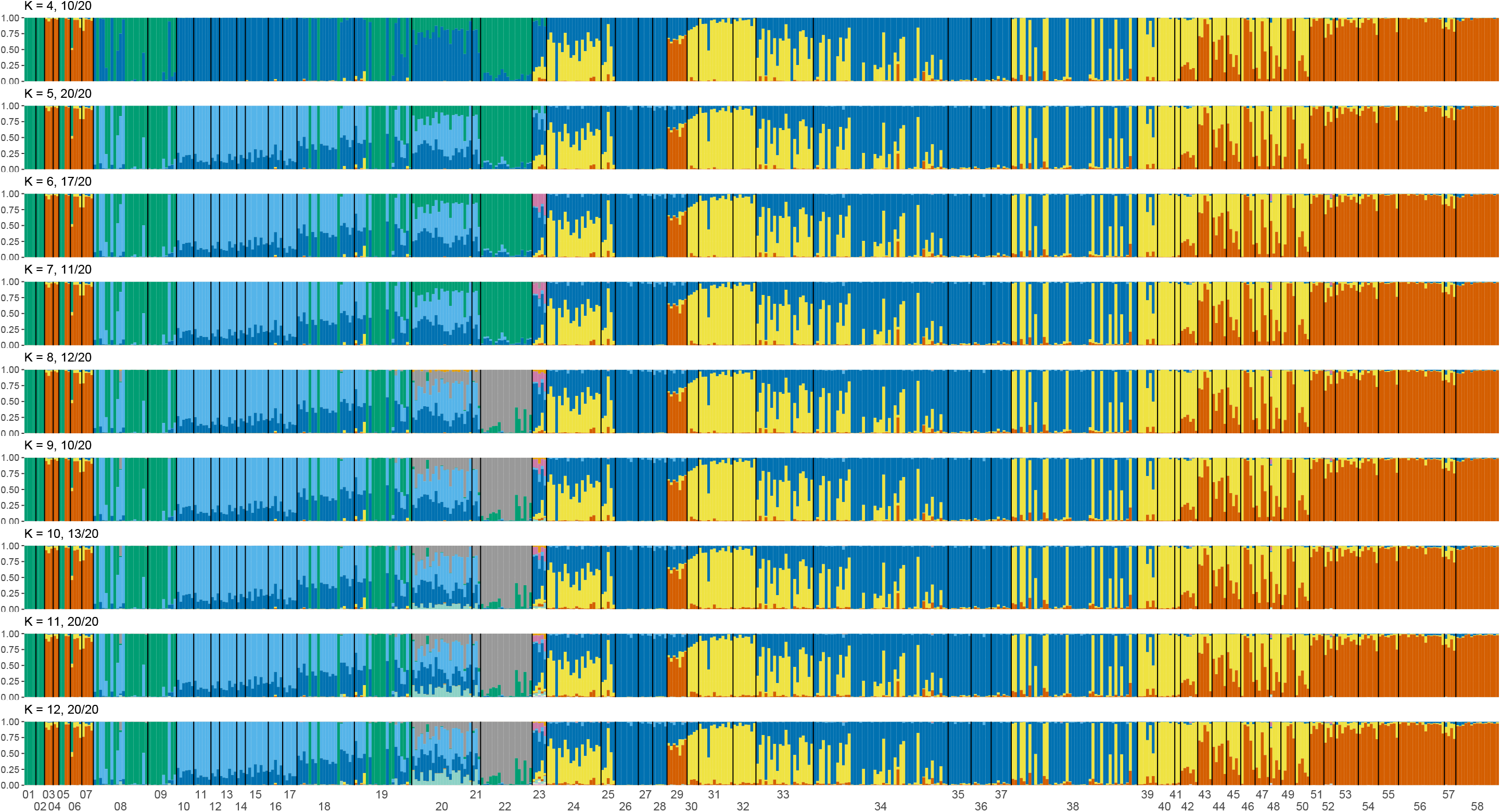
`Structure` results *K* = 4 to 12, for all populations. The major mode obtained with `Clumpak` is presented and the number of runs contributing is indicated in the panel title.

**Figure S5:**
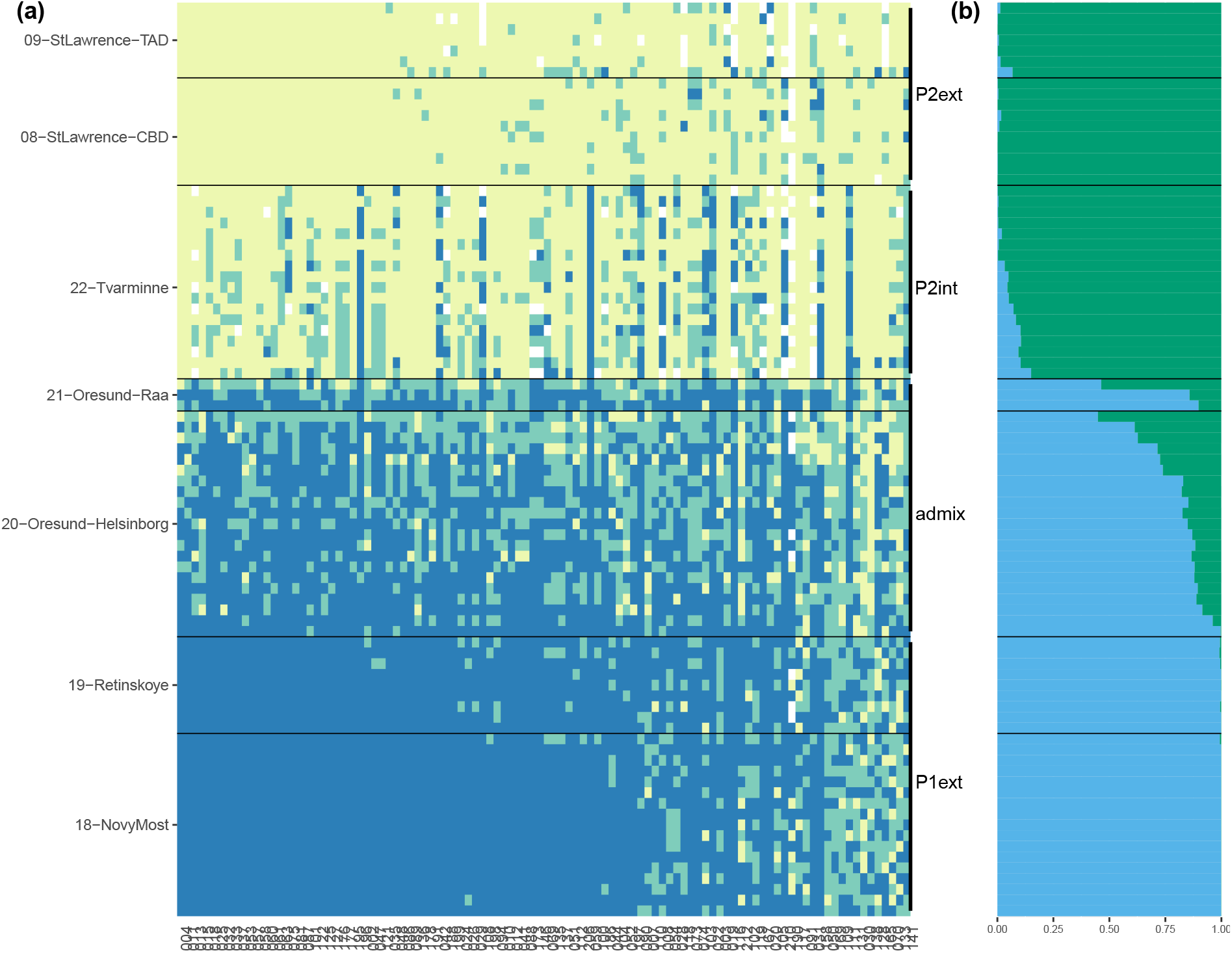
Oresund hybrid zone. (a) `introgress` plot of genotypes for each individual in line and each marker in column (AFD > 0.5). Individuals ar ordered by hybrid index in each population while markers are ordered by degree of differentiation. (b) Corresponding `Structure` ancestry compositions for each individual.

**Figure S6:**
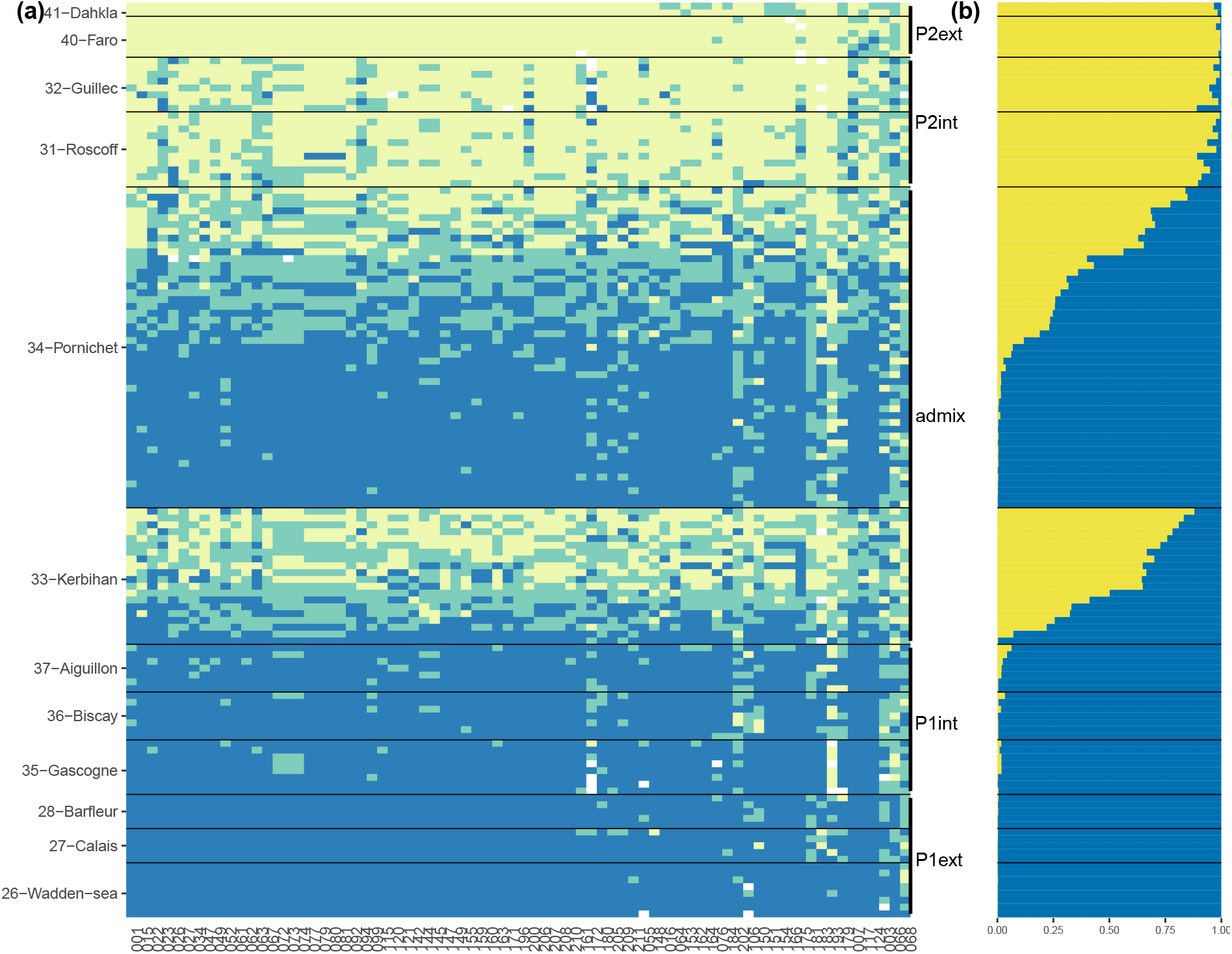
Brittany hybrid zone. (a) `introgress` plot of genotypes for each individual in line and each marker in column (AFD > 0.5). Individuals ar ordered by hybrid index in each population while markers are ordered by degree of differentiation. (b) Corresponding `Structure` ancestry compositions for each individual.

**Figure S7:**
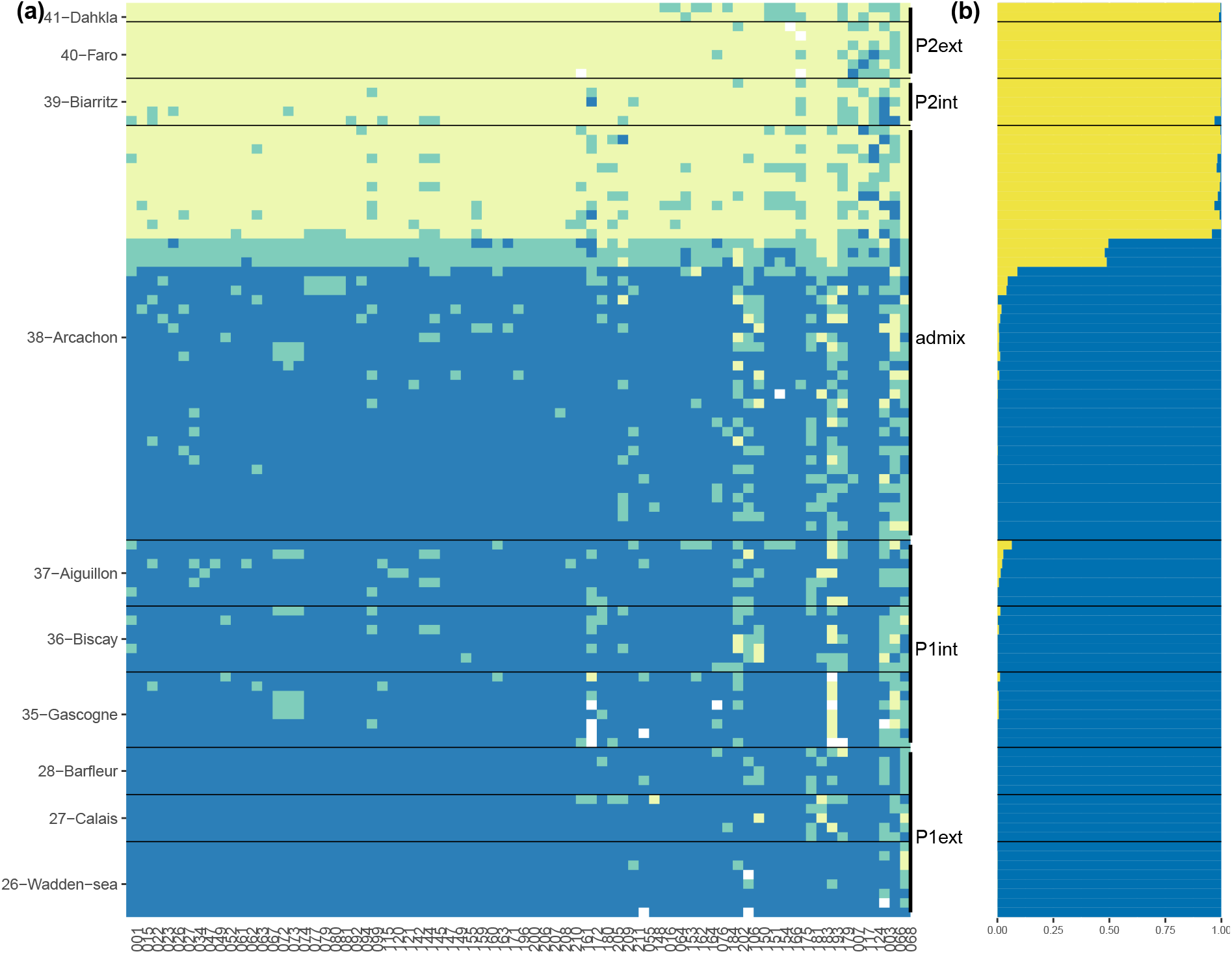
Aquitaine hybrid zone. (a) `introgress` plot of genotypes for each individual in line and each marker in column (AFD > 0.5). Individuals ar ordered by hybrid index in each population while markers are ordered by degree of differentiation. (b) Corresponding `Structure` ancestry compositions for each individual.

**Figure S8:**
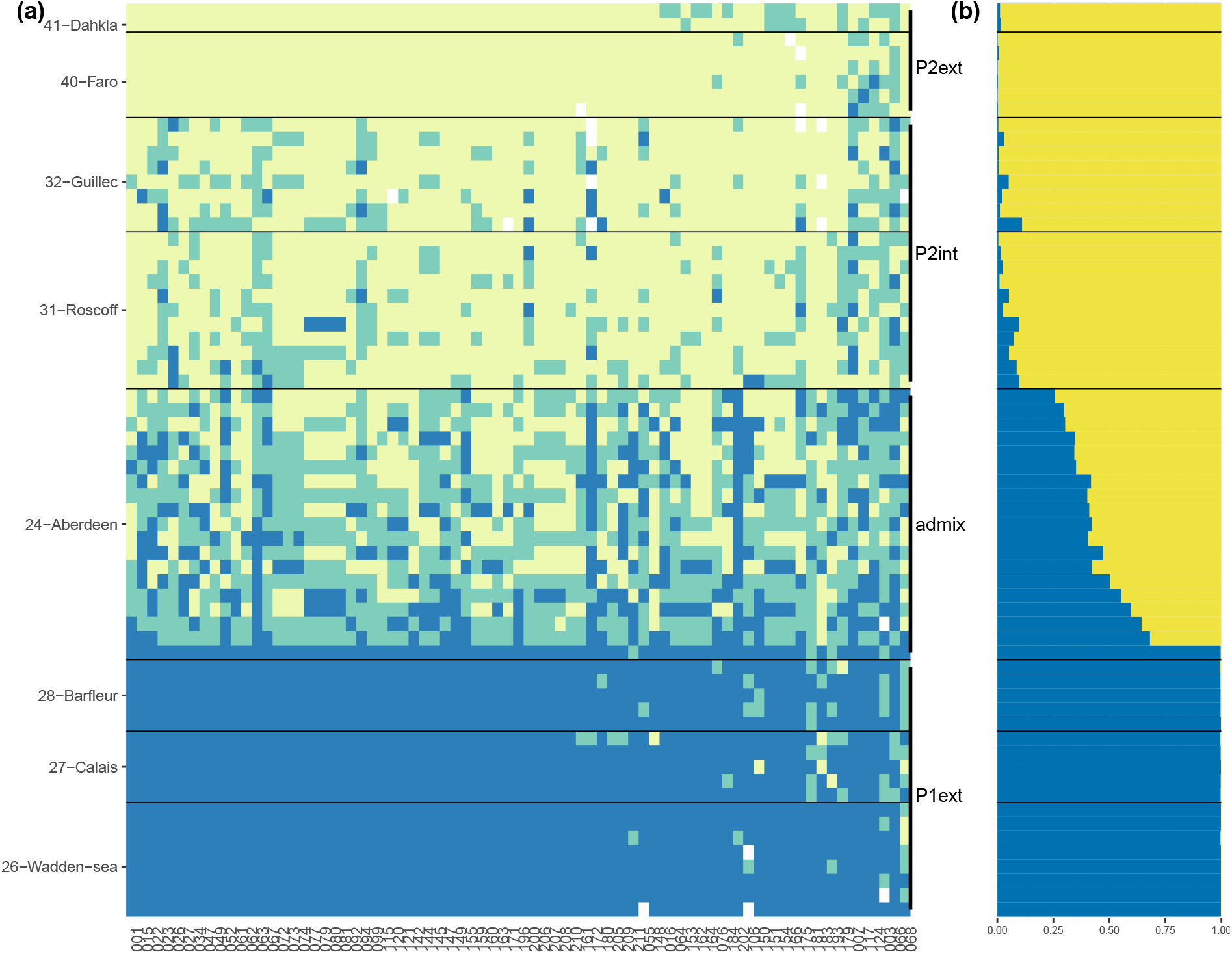
Scotland hybrid zone. (a) `introgress` plot of genotypes for each individual in line and each marker in column (AFD > 0.5). Individuals ar ordered by hybrid index in each population while markers are ordered by degree of differentiation. (b) Corresponding `Structure` ancestry compositions for each individual.

**Figure S9:**
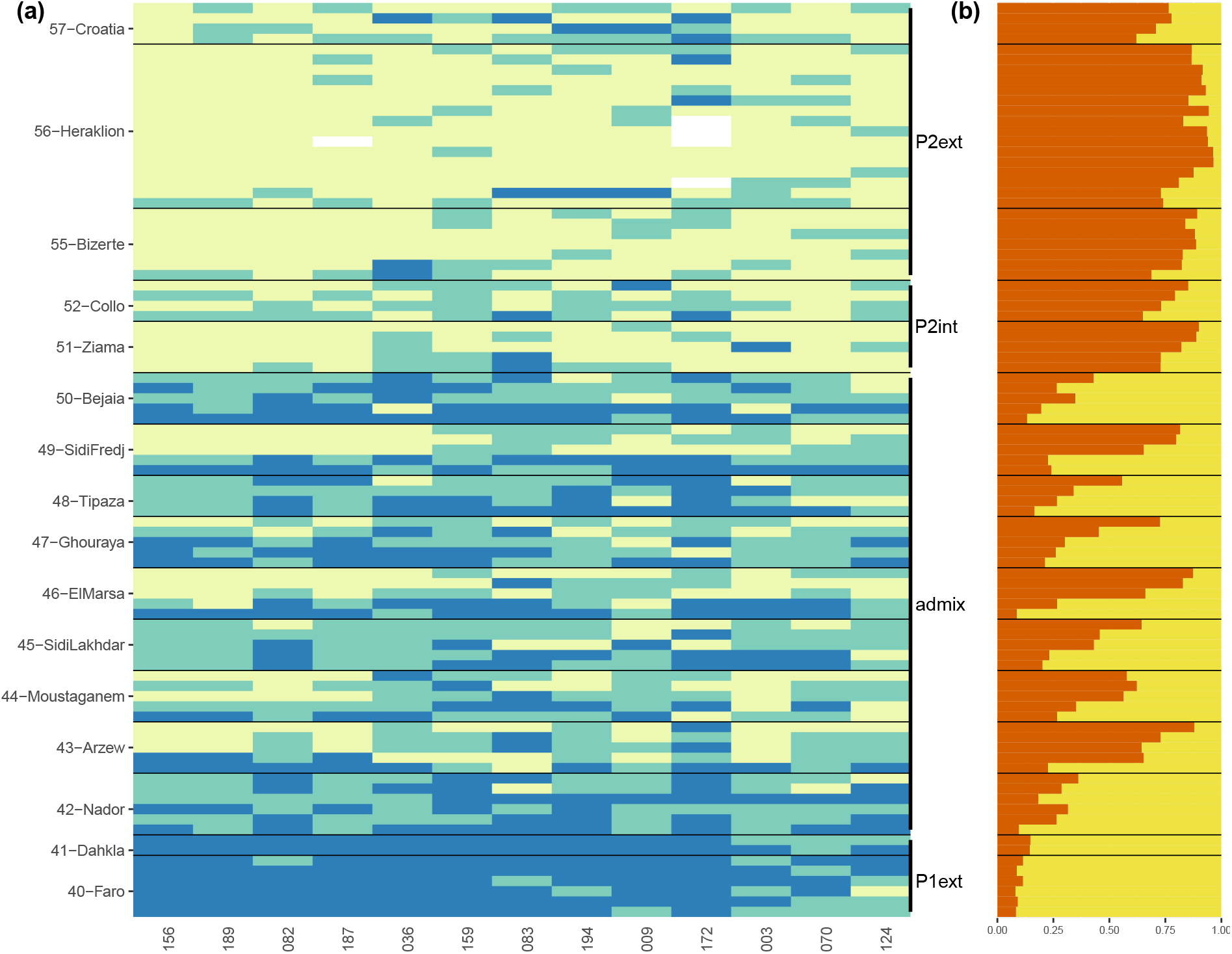
Algerian hybrid zone. (a) `introgress` plot of genotypes for each individual in line and each marker in column (AFD > 0.5). Individuals ar ordered by hybrid index in each population while markers are ordered by degree of differentiation. (b) Corresponding `Structure` ancestry compositions for each individual.

**Figure S10:**
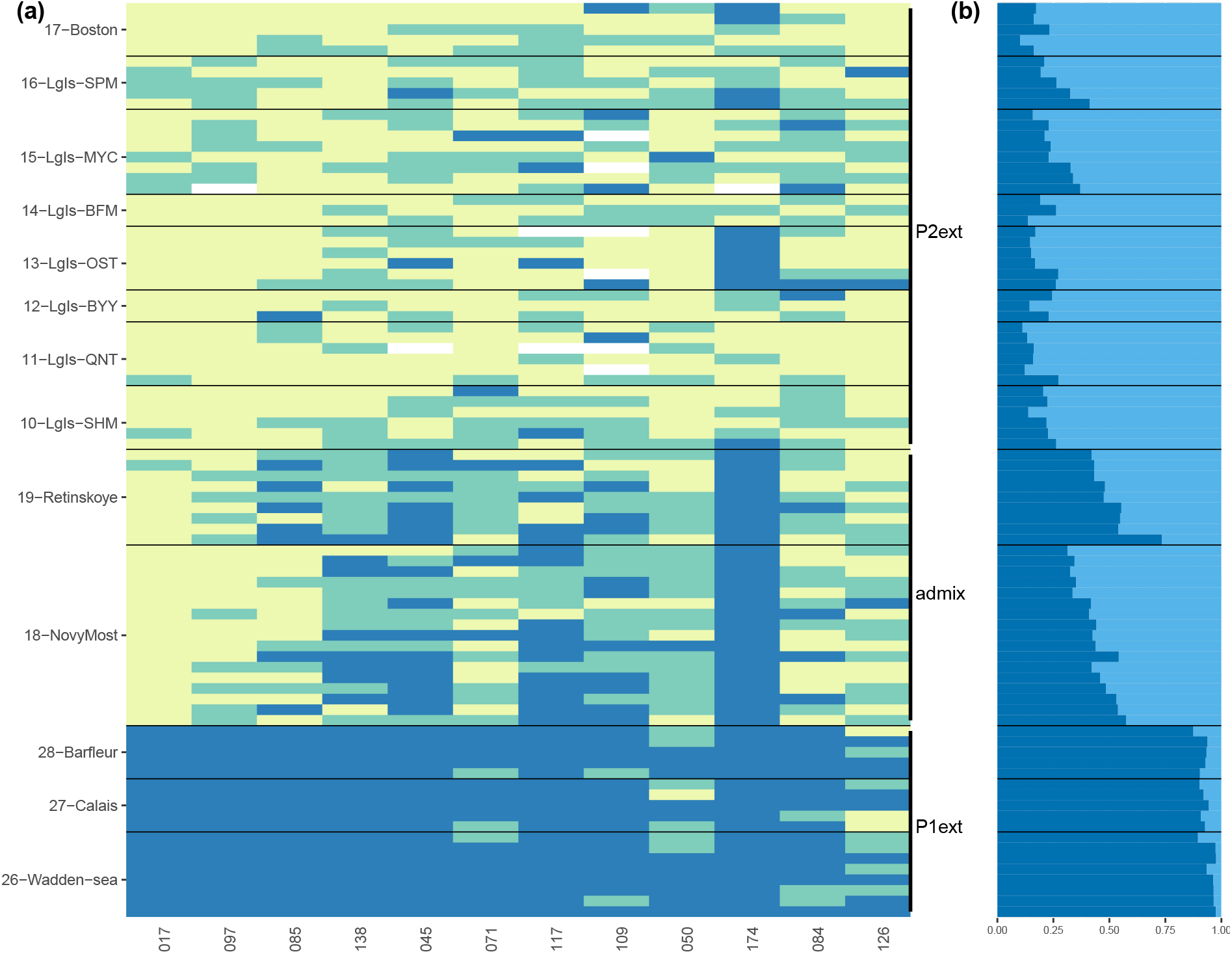
Northern European *M. edulis* hybrid zone. (a) `introgress` plot of genotypes for each individual in line and each marker in column (AFD > 0.5). Individuals ar ordered by hybrid index in each population while markers are ordered by degree of differentiation. (b) Corresponding `Structure` ancestry compositions for each individual.

